# Seeing the Forest for the trees: Assessing genetic offset predictions with Gradient Forest

**DOI:** 10.1101/2021.09.20.461151

**Authors:** Áki Jarl Láruson, Matthew C. Fitzpatrick, Stephen R. Keller, Benjamin C. Haller, Katie E. Lotterhos

## Abstract

Gradient Forest (GF) is increasingly being used to forecast climate change impacts, but remains mostly untested for this purpose. We explore its robustness to assumption violations, and relationship to measures of fitness, using SLiM simulations with explicit genome architecture and a spatial metapopulation. We evaluate measures of GF offset in: (1) a neutral model with no environmental adaptation; (2) a monogenic “population genetic” model with a single environmentally adapted locus; and (3) a polygenic “quantitative genetic” model with two adaptive traits, each adapting to a different environment. Although we found GF Offset to be broadly correlated with fitness offsets under both single locus and polygenic architectures. It could also be confounded by neutral demography, genomic architecture, and the nature of the adaptive environment. GF Offset is a promising tool, but it is important to understand its limitations and underlying assumptions, especially when used in the context of forecasting maladaptation.

## INTRODUCTION

Climate change is a growing threat to biodiversity (Urban *et al*. 2016; Nunez *et al*. 2019). Given the need to address environmental impacts on vulnerable populations, predictive models provide a means to inform conservation (Bland *et al*. 2015; Freer *et al*. 2018; Razgour *et al*. 2018). Most efforts to assess climate change impacts use species-level distribution models (Aitken *et al*. 2008; Ellis *et al*. 2012; Pacifici *et al*. 2015), but genomic data is increasingly being incorporated to provide population-level assessments (Rellstab *et al*. 2015; Hoban *et al*. 2016; Waldvogel *et al*. 2020a).

Gradient Forest (GF) is a *machine learning* algorithm that has been used to quantify and predict changes in the composition of biodiversity. GF was conceived to characterize changes in community composition (Ellis *et al*. 2012), but more recently GF has been used to identify environmental drivers of allele frequency variation, as well as to forecast the degree of maladaptation of locally adapted populations under new environments (Fitzpatrick & Keller 2015). While GF is growing in use as a predictive tool (Capblancq *et al*. 2020), neither its use in forecasting nor its application to genetic datasets have been thoroughly evaluated using “truth-known” simulations (i.e., analysis validation, *sensu* Lotterhos *et al*. 2018).

GF differs from Genotype-Environment Association (GEA) analyses (Rellstab *et al*. 2015; Hoban *et al*. 2016), which emphasizes the identification of environmentally-associated alleles, typically using linear univariate approaches (Waldvogel *et al*. 2020b). In contrast, GF fits an ensemble of regression trees using Random Forest (Breiman 2001) and then constructs *cumulative importance turnover* functions (see Table 1 for definitions) from these models by determining how well partitions distributed at numerous “split values” along each gradient explain changes in allele frequencies on either side of a split. These cumulative importance curves are generated for each fitted response (e.g., a single nucleotide polymorphism (SNP), or a single species count) and each environmental predictor, which are weighed and combined to produce an aggregate cumulative importance curve for the genome (or a community of species) along each significant predictor. The slope of a SNP-level cumulative importance curve should indicate the rate of change in the allele frequency across the environmental gradient, but this remains untested.

**Table 1.** Terminology

| Terms | Description |
| --- | --- |
| Causal Environment | Environmental variable that determines the optimal phenotype |
| CG Fitness<sub>*m,n*</sub> | Common Garden (CG) Fitness averaged across all individuals from an *m* source location in an *n* transplant (common garden) location |
| Cumulative Importance | The cumulative sum of “split values” from the fitted GF model. |
| Fitness offset | The difference in fitness experienced by moving an organism from its home environment to a foreign one. |
| $F_{ST \text{ causal}}$ | Genetic differentiation between the source population and transplant (common garden) population at loci with alleles that have causal effects on the phenotype |
| $F_{ST \text{ genome}}$ | Genetic differentiation between the source population and transplant (common garden) population across the genome |
| GF Offset | The euclidean distance between the cumulative importance output by Gradient Forests at one multivariate environment and another multivariate environment |
| Local Adaptation | The difference between the fitness of populations in sympatry and allopatry |
| Machine Learning Algorithm | An algorithm that builds a model based on a subsample of the data, in order to make predictions for an expanded dataset |
| Relative Fitness in common garden | The relative probability that an individual with a given genotype would be a parent to offspring in the next generation in the common garden environment, given the causal alleles it possesses and the functions relating alleles to phenotype and phenotype to fitness |
| Turnover | A change in allele frequency across an environmental gradient |
| Vulnerability | The extent to which an organism is susceptible to or unable to cope with climate change and includes the magnitude and rate of exposure to climate change, sensitivity to that exposure, and the ability to cope with climate-related changes through adaptive capacity. See Foden *et al.* 2019. |

In addition to providing inference regarding the nature of allele frequency change along spatial environmental gradients, GF has been proposed as a method to predict the frequency change in locally adaptive alleles needed to maintain current levels of adaptation following a change in environment (Fitzpatrick & Keller 2015; Fitzpatrick *et al*. 2018; Capblancq *et al*. 2020). In essence, GF’s turnover functions provide a means to transform (i.e., rescale) environmental predictors from their original units (e.g., °C, mm) into common units of cumulative importance. The transformed predictors can then be used to calculate expected genetic differences as the Euclidean distance between populations in time and/or space (Gougherty *et al*. 2021), a distance referred to as “genetic offset” by Fitzpatrick and Keller (2015) and as “genomic vulnerability” by Bay *et al*. (2018) and Ruegg *et al*. (2018). We refer to this distance as “GF Offset” to emphasize its derivation from GF and we suggest the term “genomic vulnerability” should be avoided given that (1) it does not fit established definitions of climate change *vulnerability* (Foden *et al*. 2019) and (2) it is not clear to what extent GF Offset represents vulnerability, however defined.

Several questions regarding the use of GF Offset as a metric of maladaptation (e.g., assuming increased GF Offset corresponds to increased *fitness offset*) remain unanswered, including how it is affected by neutral demography. For example, changes in allele frequencies could reflect simple genetic drift rather than adaptive signals (Rellstab *et al*. 2015; Hoban *et al*. 2016; Fitzpatrick *et al*. 2018; Borrell *et al*. 2020). Smaller populations will tend to exhibit greater signatures of drift than larger populations (Wright 1929; Buri 1956; Helgason *et al*. 2003) and potentially greater allele frequency turnover in regions with smaller population sizes, and less turnover in regions with large population sizes. If these gradients in population structure are aligned with environmental gradients, they will be reflected in the fitted cumulative importance curves from GF (i.e., steeper slopes where allele frequency turnover is high, flatter slopes where turnover is low) and therefore could artificially inflate predicted GF Offsets.

In addition, a complexity inherent to interpretations of GF Offset in the context of forecasting maladaptation is that it is a multivariate distance of allele frequencies from a presumed optimum, meaning that regardless of the direction of change (increase or decrease) in allele frequencies across a gradient, GF Offset will always be positive. The underlying assumptions being that a population already occupies its adaptive optimum when sampled and therefore any change in allele frequency composition will reduce fitness. However, because fitness could *decrease* or *increase* in response to environmental change, it is unclear how GF Offset actually relates to fitness, especially when contrasted with other better known distance metrics, such as *F_ST_*. To evaluate GF Offset as a potential measure of fitness offset, we sought to address the following questions:

Q1) *Variation in N*. What effect do neutral processes, operating on a cline in population size across an environmental gradient, have on GF Offset? We tested the hypothesis that a decrease in population size would result in an increased GF Offset, due to increased genetic drift operating in small populations.
Q2) *Relationship between GF Offset and fitness offset*. Given equal deme sizes in a metapopulation, how well does GF Offset predict changes in fitness when populations experience an immediate environmental change (i.e., with no associated evolutionary change)? We tested the hypothesis that GF Offset is inversely related to fitness, by conducting *in silico* common garden experiments under monogenic and polygenic architectures.
Q3) *GF Offset versus other measures of offset*. Given equal deme sizes in a metapopulation, how does GF Offset perform relative to environmental distance or *F_ST_*? We tested the hypothesis that GF Offset outperforms both environmental distance and genetic distance, because GF appropriately weights and scales the environmental gradients to reflect their genetic importance.

We tested the performance of GF and other offset measures in their ability to predict fitness of genotypes when transplanted to common gardens (avoiding the confounding longer-term dynamics of dispersal and gene flow) across the species range *in silico*. Using SLiM (Haller & Messer 2019), we simulated different genome architures that underlie a phenotype, and different relationships between the phenotypic optimum and a single environmental variable. We then evaluated GF Offset from three scenarios: (1) a neutral model with clinal population size across the environment; (2) a monogenic “population genetic” model with adaptation to a single environment; and (3) a polygenic “quantitative genetic” model with two environmentally adaptive traits, each responding to a different environmental gradient. Overall we find that GF Offset was strongly correlated with fitness offset, but that there are multiple sources of confounding effects.

## MATERIALS & METHODS

### Demography and genetic architecture

We generated simulations using SLiM v3.4. Ten thousand individuals (*N*) were split across a metapopulation consisting of 100 demes arranged in a 10×10 connectivity matrix (**Fig S1**- [supp](https://docs.google.com/document/d/1oCI5_FIs9DvaAkSVCtFFALqwBvnscTNnUXLB3xxX1Is/edit#heading=h.bujj4m9yp26c)). Each deme (*D_x,y_*) was assigned at least one environmental value that could vary over generational time (*E_j,t_*, where *j* is a deme, and *t* is the generation). When two environmental variables were considered, they are referred to as Environment1 and Environment2 (*E1_j,t_* & *E2_j,t_*). Symmetric migration was simulated between adjacent demes at a per-generation migration rate (*m*), and each deme contained equal proportions of males and females, with bi-parental mating producing the subsequent generation.

Ten genomic linkage groups each containing 50K sites were simulated in each individual, for a total of 500K sites per haploid copy of the diploid genome. The neutral mutation rate (*μ*) was10^−7^(a metapopulation-scaled mutation rate *N_T_* * *μ* of 0.001), and a base recombination rate (*r*) of 10^−5^(*N_T_* * *r* = 0.1) was used to approximate a distance of 50 cM per linkage group. The population-scaled mutation and recombination rates were chosen to approximate the resolution that would be observed from sampling SNPs from a larger genome, *i.e*., allowing SNPs 50K bases away to be unlinked, while allowing for signatures of linkage to arise within each linkage group (Lotterhos 2019). All simulation parameters are listed in Table 2.

**Table 2.** Model Parameters

| Parameter | Value | Description |
| --- | --- | --- |
| N | 10,000 | Total population size of the meta-population |
| n | 100 | Individual population size |
| m | 0.05/0.2 | Per-generation migration rate (single locus/multi locus) |
| $\mu$ | $1 \times 10^{-7}$ | Neutral mutation rate |
| r | $1 \times 10^{-5}$ | Recombination rate |
| d | 0.45 | Fitness slope parameter (single locus model) |
| $\mu_{\text{QTN}}$ | $2.5 \times 10^{-6}$ | QTN mutation rate (multi locus models) |
| $\sigma_{\text{QTN}}$ | 0.1 | Variation in QTN effect sizes (multi locus models) |
| $\sigma_{\text{S}}$ | 4.0 | Strength of stabilizing selection for first 1000 generations (multi locus models) |
| $\sigma_{\text{K}}$ | 1.25/4.0 | Strength of stabilizing selection after 2000 generations (multi locus cases 1,2,3/multi locus case 4 for second trait) |

Simulations were output as tree-sequence files (Haller & Messer 2019; Haller *et al*. 2019) following a burn-in period and a period of stable environmental values (the length of these periods is explained below). The software packages `msprime` and `pyslim` were used to prepend a simulation of neutrally coalesced ancestry onto the SLiM-generated file to guarantee every site was fully coalesced (referred to as “recapitation”), and neutral variants were then overlaid on that recapitated file (Haller *et al*. 2019). We used `vcftools` to filter for minor allele frequencies above 0.01, which is on the lower range of MAF filtering criteria in genomics studies (Danecek *et al*. 2011; Byrne *et al*. 2013; Linck & Battey 2019) but ensures that in our multilocus simulations we included more local causal alleles in our analyses. Out of the 10,000 individuals simulated, 10 individuals from each deme (for a total of 1,000 individuals) were randomly selected for downstream analysis.

### Thought experiments

To elucidate what drives the shape of the cumulative importance function, we created five allele frequency patterns across a gradient representing an environmental variable and analyzed them in GF: (1) different sampling schemes of a steep allele frequency cline along one or multiple environmental variables, (2) different slopes of allele frequency clines along an environmental variable, (3) different non-monotonic relationships between allele frequency and an environmental variable, (4) a comparison of linear and non-monotonic allele frequency relationships with an environmental variable, and (5) a comparison of a linear allele frequency relationship with an environmental variable and the same relationship with noise added by sampling from a normal distribution with 0 mean and variance of 0.1.

### Variation in N

#### Wright–Fisher neutral model

To test the hypothesis that GF Offset could be influenced by variation in genetic drift, we simulated a linear environmental gradient with values from −1 to 1, in three neutral scenarios: equal deme size (*N_d_*); increasing *N_d_*; and decreasing N_d_ (**Fig 1**). In the equal *N_d_* scenario, each deme contained 100 individuals. In order to maintain a consistent *N_T_* for the increasing and decreasing deme size scenarios, the sum of *N_d_* within each row of the metapopulation grid was made to equal 1,000 individuals (e.g., increasing *N_d_* scenario D_1y-10y_: 10, 10, 50, 50, 95, 95, 145, 145, 200, 200; the decreasing *N_d_* scenario was the opposite). Each scenario was replicated ten times.

**Figure 1.**
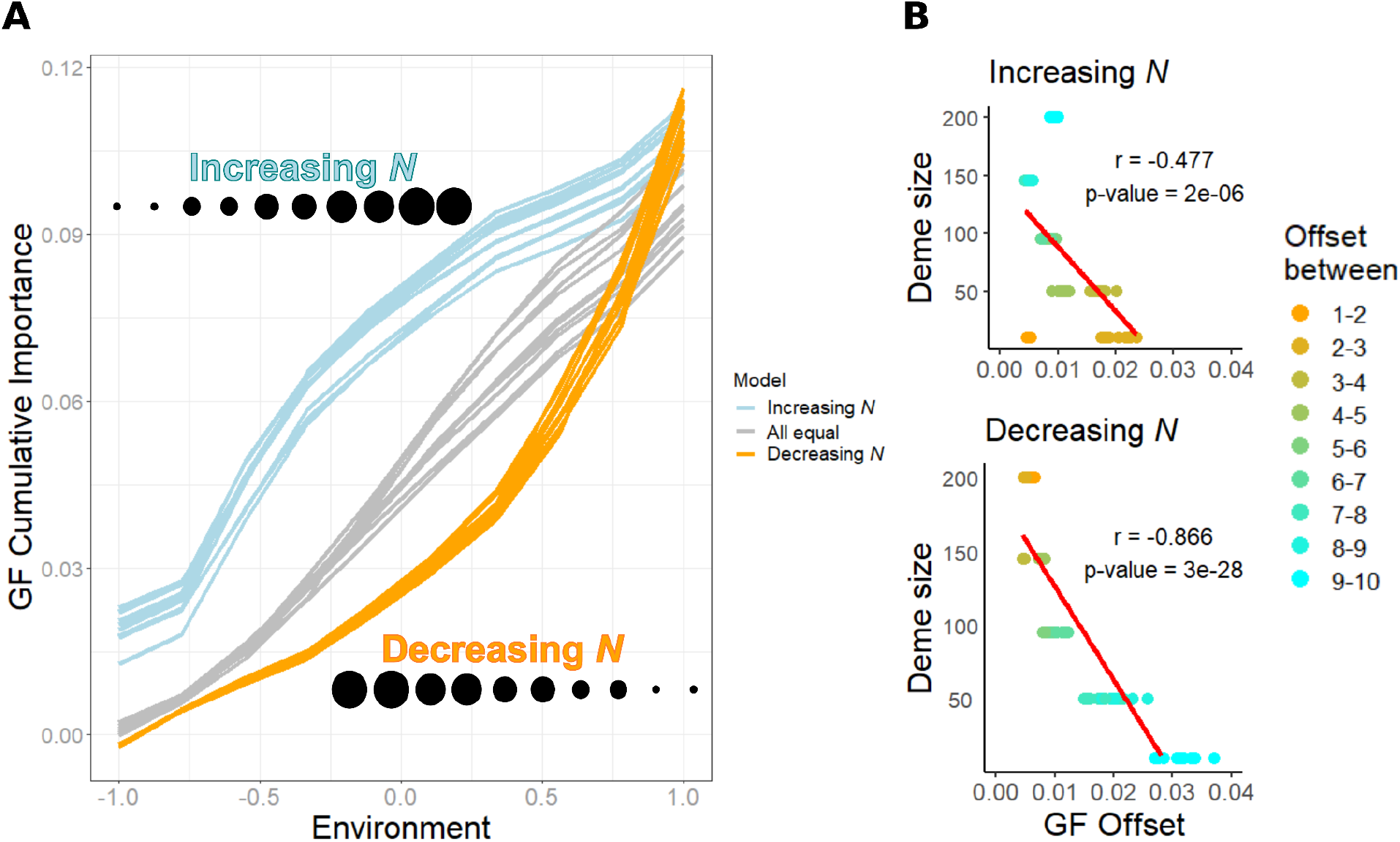
Results of neutral simulations, where no selective pressure is imposed by the underlying environmental clines. **A**. The relationship between GF Offset and Environment across increasing, decreasing, and equal deme sizes across the environmental gradient. GF Offset increased comparatively more when deme sizes were small. **B**. A strong negative linear Pearson’s correlation between the GF Offset and deme size in ten replicates, regardless of the direction of the population gradient. Numbers 1 through 10 in the legend represent the columns in the metapopulation matrix (see Figure MetaPop figure).

#### Q1: Effect of N_d_ on GF Offset

GF analysis, implemented in R (R Core Team 2021) in the ‘gradientForest’ package (Ellis *et al*. 2012), was performed using filtered allele frequencies and environmental values sampled as described for N_d_ above. We measured the GF Offset of each deme based on an adjacent environmental shift. We tested the null hypothesis of no relationship between *N_d_* and the GF Offset with Pearson’s correlation coefficient in R. If GF Offset is not affected by genetic drift, then this correlation should equal 0. However, if higher drift at one end of an environmental cline results in more allele frequency turnover (and higher GF offsets for those populations), then the correlation between *N_d_* and GF Offset should be negative.

### Q2: Relationship between GF Offset and fitness offset

Our goal was to use these simulations to assess the predictive potential of GF Offset for a straightforward demography, in which population size was equal for all demes. These simulations evolved a simple isolation-by-distance population structure. We evaluated the relationship between GF Offset and mean deme fitness, after individuals from that deme were transplanted into another environment in the simulation, for both monogenic and polygenic scenarios. In order to reduce computation load, we explore best-case scenarios for GF (e.g., where there is strong environmentally driven local adaptation), migration values were chosen which gave high levels of local adaptation, such that the average deme fitness was approximately 15–25% greater in the home environment than in allopatric environments.

#### Relationship between environment and fitness

##### Single-locus single-environment population genetic simulations

In our “population genetic” model of environmental adaptation, a single allele had a linear relationship with fitness across an environmental cline. To avoid a scenario where maladapted individuals persist at the range edge, we modeled individual fitness for each genotype as a function of the environment, with the ancestral allele (*a*) considered to be antagonistically pleiotropic (*sensu* Savolainen *et al*. 2013) to the emerging derived allele (*A*). See [supplement](https://docs.google.com/document/d/1oCI5_FIs9DvaAkSVCtFFALqwBvnscTNnUXLB3xxX1Is/edit#) S1 for more details.

##### Multilocus two-trait two-environment quantitative genetic model

In our “quantitative genetic” two-trait, two-environment model of environmental adaptation, QTNs were allowed to arise at a rate of *μ_QTN_* = *μ/4* = 2.5×10^-6^ (Table 2) across 9 of the 10 linkage groups in the genome, with the final linkage group maintained as a neutral genomic reference. We assumed a quantitative genetic model where alleles contributed additively to two distinct phenotypes for each individual *i* in deme *j* (*P*_1,*ij*_ and *P*_2,*ij*_) with no dominance. When an ancestral allele mutated, the bivariate effect size of the derived allele was drawn from a multivariate normal distribution with a standard deviation of *σ*_QTN_ = 0.1 (without covariance) for both traits, which gives flexibility for mutations to evolve with effects on one or both traits (i.e., pleiotropy). For each deme the phenotypic optimum simply equaled the environmental value.

The relative fitness of individual *i* in deme *j* (*ω_ij_*) was based on how far their *P*_1,*ij*_ and *P*_2,*ij*_ phenotypes fell from the bivariate optimum in that patch (Θ_1jt_ and Θ_2*jt*_) using a multivariate normal distribution with standard deviation *σ_k_* to represent the strength of stabilizing selection in each deme:

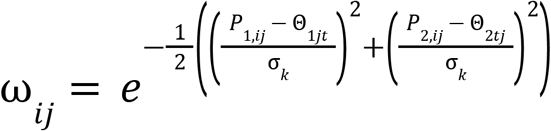

All relative fitness values were normalized by *ω*_max_, where *P_ij_* = Θ_*jt*_. We ran a cumulative burn-in period of 3000 generations, consisting of a 1000 generation homogeneous initial burn-in, followed by a 1000 generation transition burn-in (see [supplement](https://docs.google.com/document/u/0/d/1oCI5_FIs9DvaAkSVCtFFALqwBvnscTNnUXLB3xxX1Is/edit) **S1**).

#### Local adaptation and migration

An elevated migration rate (*m* = 0.2) was chosen to allow a high degree of local adaptation to quickly arise across the metapopulation. We measured the mean local adaptation across each simulation as the difference between mean sympatric fitness (*ω*_S_) and mean allopatric fitness (*ω*_A_) (Blanquart *et al*. 2013). Mean sympatric fitness was quantified as the average value along the diagonal of the common-garden fitness matrix, while mean allopatric fitness was quantified as the average of all of the off-diagonal values.

##### Case scenarios

Four cases were considered within the multilocus model (**Fig 3**). Except as noted, the post-burn-in optima ranged from −1 to 1 for both environments, and the strength of selection equaled 1.25. Case 1 simulates simple linear clines, in which genetic distance is linearly related to environmental distance. Two orthogonal environmental clines were simulated: environment 1 varying left-to-right, environment 2 varying bottom-to-top (**Fig 3 Case 1**). Case 2 simulates a situation in which geographic distance does not relate linearly to environmental distance. Here we simulated two non-monotonic environments, with the two orthogonal environments increasing from one edge to the middle of the metapopulation, and then decreasing again to the opposite edge. The 4 corners of the landscape thus had the same environment, but were geographically distant (**Fig 3 Case 2**). In Case 3, we sought to understand the effect of the slope of the environmental gradient on GF Offset. Case 3 was set up identically to Case 1 except that the optima for environment 2 were narrowed, ranging from −0.25 to 0.25. This was done to produce a narrower phenotypic range in one environment (which should evolve weaker clines in allele frequency), versus another, without changing the strength of selection (**Fig 3 Case 3**). In Case 4, we sought to determine whether the strength of selection affects GF Offset. Case 4 was set up identically to Case 1 except that environment 2 had weaker stabilizing selection, *σ*_k_ = 4 for ϴ_2_ to produce less overall local adaptation, and more additive genetic variation, for trait 2 than produced by Case 1 (**Fig 3 Case 4**). Each Case was replicated ten times.

#### Fitness assessment in a Common Garden - CG Fitness

In order to represent the fitness effects of rapid climate change, we implemented a reciprocal transplant fitness assessment in our single- and multi-locus simulations. The mean relative fitness of individuals from a home location *i* (*D*_x,y_) in a transplant “common garden” location *j* (*D*_x’,y’_) was calculated for each deme across all contemporary environments in the simulation, resulting in a 100×100 matrix of pairwise fitness comparisons. For each common garden *j*, we could then assess how well the GF Offset between *i* and *j* predicted the relative fitness of individuals from *i* when transplanted to *j* (see Methods: Evaluating GF Offset). We refer to this fitness measure as *CG Fitness* (common garden fitness).

#### Q3: Offset measures

In order to assess the performance of GF Offset in predicting CG Fitness relative to other distance metrics, we also calculated *F*_ST_ values and environmental distances. Pairwise Weir–Cockerham *F*_ST_ values were calculated for every combination of demes at the end of each simulation after filtering loci for MAF: one set of such values using all alleles (*F*_ST Genome_), another set using only the QTN alleles (*F*_ST Causal_). Environmental distances were calculated using both *n*-dimensional Euclidean (*E_D_*, Cauchy 1882) and Mahalanobis (*M_D_*, Mahalanobis 1930) distances between all pairwise demes.

We also evaluated the effect of adding non-causal environmental variables into the offset calculations by simulating an additional 12 environmental gradients. Two of these gradients were correlated with the two causal gradients, derived by adding random draws from a univariate normal (μ = 0, σ = 1.3, to allow for a Pearson’s correlation between 0.4-0.5). The other ten gradients were drawn from a multivariate normal distribution using a covariance matrix generated by sampling the correlation among variables from a uniform distribution (clusterGeneration package v1.3.4), which gave them a correlation structure similar to that observed in climate data. Distances were calculated using all environmental variables (“total environmental distance”, *M_D-all_, E_D-all_*, 14 variables), and using just the causal environments (“causal environmental distance”, *M_D-causal_, E_D-causal_*, 2 variables).

GF outputs an individual cumulative importance curve for each locus and a weighted aggregate function across all loci, for each important environmental variable. GF Offset is defined as the Euclidean distance between two locations *A* and *B* in the rescaled environmental space (or *spaces* over time) obtained by applying the fitted GF model to the environmental predictors. Note that the rescaling and calculation of GF Offset can be performed using the individual cumulative importance curves (if the goal is to calculate the offset for a single locus, e.g., (Keller *et al*. 2018)), but most applications to date have used the aggregate cumulative importance curves and therefore calculated a multi-locus offset:

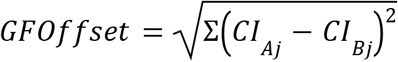

where *CI_Aj_* is the multivariate cumulative importance calculated at point *A* for environment *j*, and *CI_Bj_* is the same variable calculated at point *B*.

For each common-garden site, we measured the correlation (Spearman’s rho) between each offset prediction and the CG Fitness for that site. It became apparent that the relationship between GF Offset and CG Fitness could depend on the common garden location, so explicit comparisons to measures of CG Fitness were performed with two subsets of the metapopulation, an Edge and a Core set of demes (**Fig S1** - [supp](https://docs.google.com/document/d/1oCI5_FIs9DvaAkSVCtFFALqwBvnscTNnUXLB3xxX1Is/edit#heading=h.bujj4m9yp26c)).

## RESULTS

### Thought experiments

To explore the behavior of GF, we created different relationships between allele frequency and an environmental cline and used these to evaluate the relationships between the rate of allele frequency turnover and total amount of CI, the shape of the CI curve, and the *R^2^* value. GF produces the same nonlinear cumulative importance curves for linear clines, in which the rate of turnover was highest near the middle of the cline and low elsewhere, regardless of the slope between allele frequency and the environment (**Fig 2**, “steep”, “reverse”, and “shallow” clines). Non-monotonic allele frequency patterns also produced a nonlinear cumulative importance curve, the shape of which matched the rate of turnover in allele frequencies, namely for values of the environment where there is rapid turnover in allele frequencies, the slope of the curve is high and in contrast, where there is no allele frequency turnover in the non-monotonic case, there is also no increase in the cumulative importance (**Fig 2**, “non-monotonic”). Note how in the “non-monotonic case” the allele frequency is the same at extremes of the environmental gradient, but (given the assumption of monotonicity in the construction of cumulative importance curves) a deme would be predicted to have a nonzero GF Offset for an environmental shift from −1 to 1 for that locus. See supplement **S2** for results from a more comprehensive set of thought experiments, which illustrate how the number of sampled populations and random error influence cumulative importance curves (and thus GF Offset).

**Figure 2.**
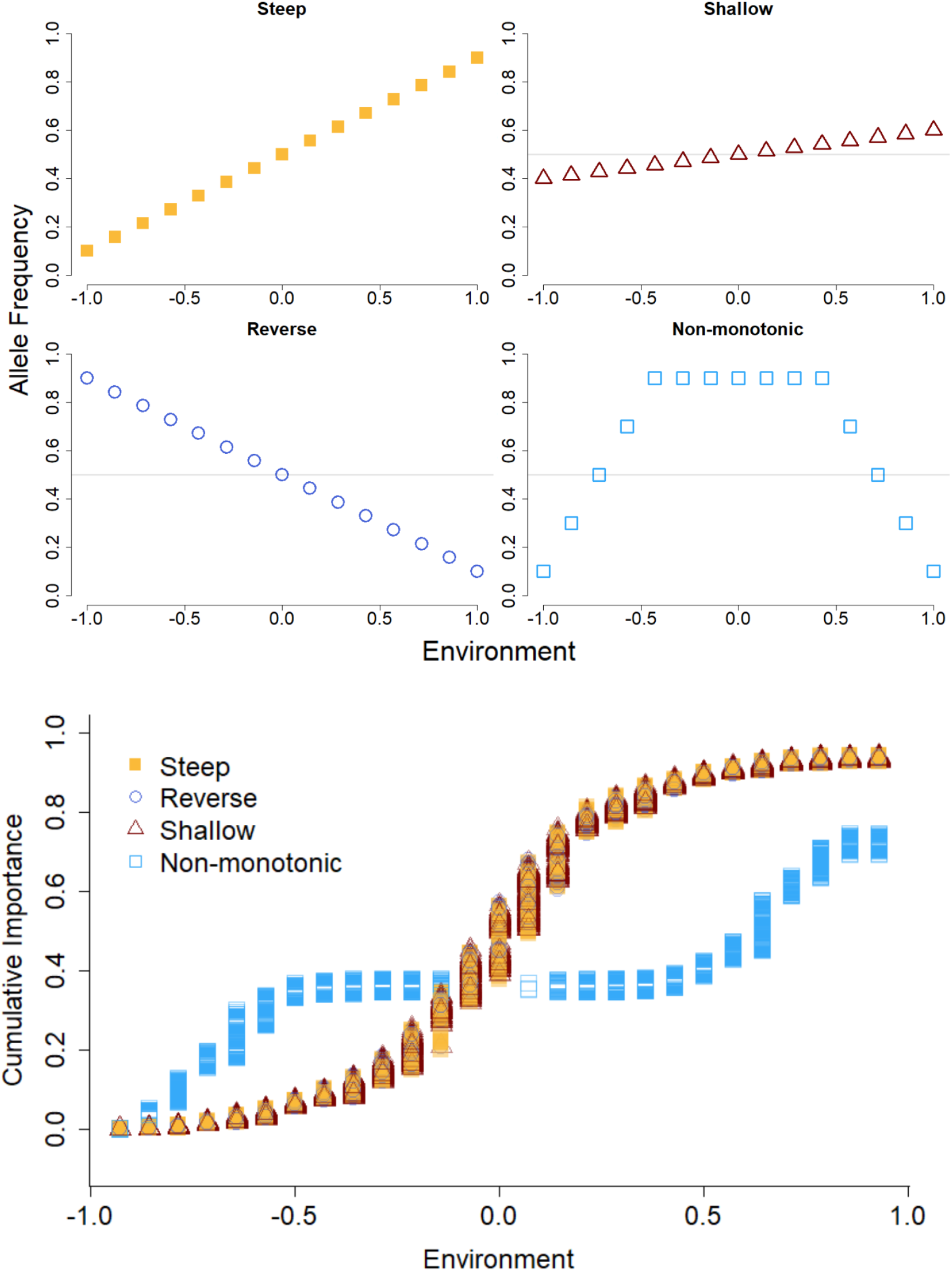
Different allele frequency gradients (steep, shallow, reversed, and non-monotonic) with respect to an environmental gradient, and their corresponding cumulative importance curve produced by Gradient Forest.

**Figure 3.**
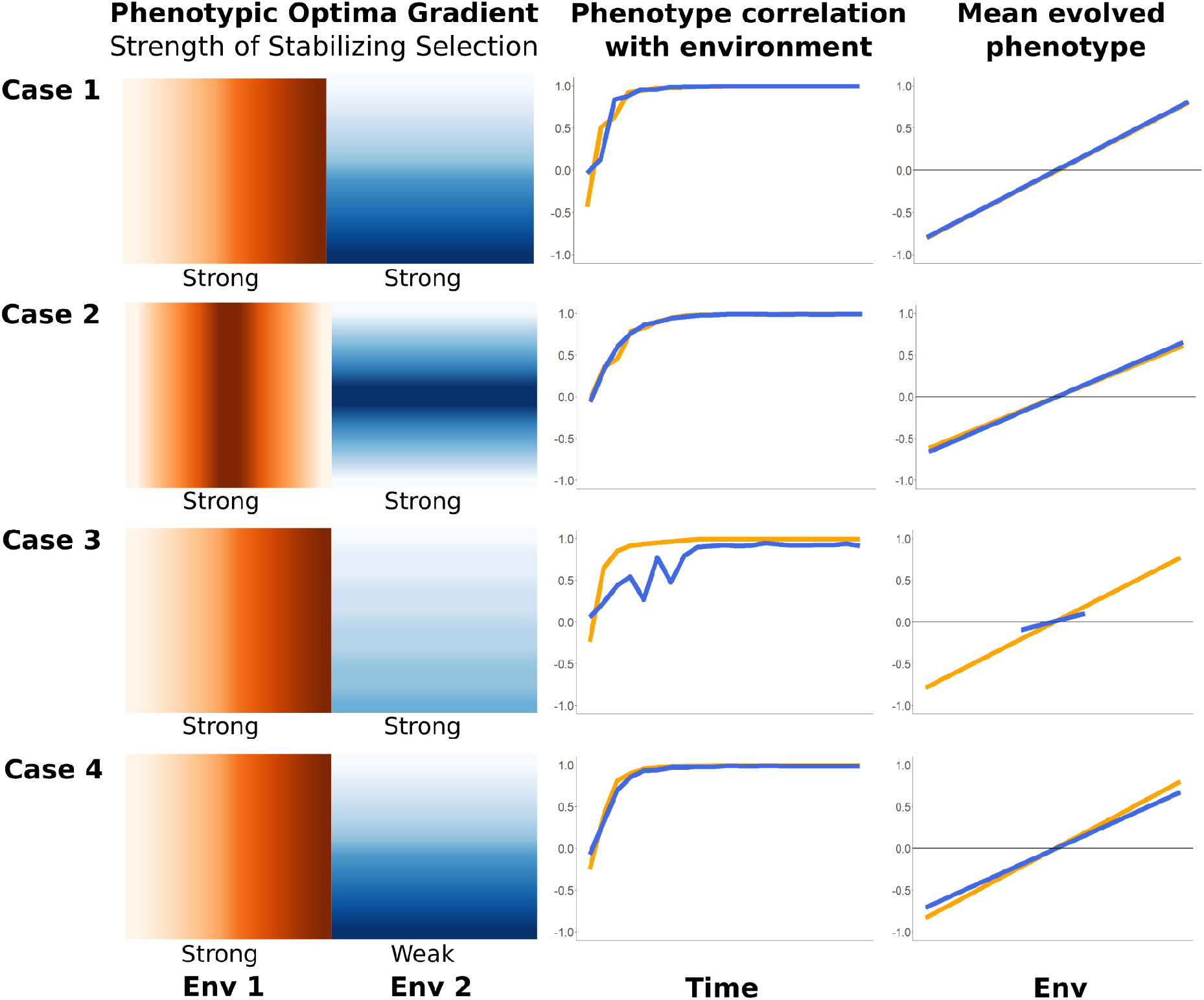
Visualization of the four multilocus case studies. Rows summarize the four key features of each case: the clines of the two phenotypic optima shown as color gradients, with the strength of stabilizing selection listed below each gradient (left column); the correlation of each evolved phenotype to its respective environment over time (middle column); and the mean value of each evolved phenotype at the end of the simulation plotted against its home environment (right column).

### Variation in N

When deme sizes across the environment were uniform, the cumulative importance increased linearly in all replicates (**Fig 1A**). In contrast, when deme sizes increased or decreased along the environmental gradient, the rate of increase in the cumulative importance was steeper at smaller deme sizes and less steep at larger deme sizes (**Fig 1**). In all replicates, GF Offset was negatively correlated with deme size (**Fig 1B**).

### Relationship between GF Offset and fitness offset

#### Single locus case

When a single locus of large effect drives environmental adaptation, GF readily identified the environment driving the clinal pattern (**Fig S2** - [supp](https://docs.google.com/document/d/1oCI5_FIs9DvaAkSVCtFFALqwBvnscTNnUXLB3xxX1Is/edit#)). The correlation between GF Offset and CG Fitness was negatively correlated in common gardens at the edges of the range. In the range center there was also significant negative correlation, but the slope was almost flat, indicating that populations had similar fitness in that garden despite a range of offset values. (**Fig S3** - [supp](https://docs.google.com/document/d/1oCI5_FIs9DvaAkSVCtFFALqwBvnscTNnUXLB3xxX1Is/edit#))

#### Multilocus cases

All simulated cases produced high levels of local adaptation (**Fig S4** - [supp](https://docs.google.com/document/d/1oCI5_FIs9DvaAkSVCtFFALqwBvnscTNnUXLB3xxX1Is/edit#)), with Case 3 having the lowest (0.169 - on average an individual was 16.9% more fit in home environment) and Case 1 having the highest (0.288). In the cases with 2 linear causal environmental clines (Cases 1, 3, & 4), GF was able to identify the causal environment driving adaptation (**Fig S5** - [supp](https://docs.google.com/document/d/1oCI5_FIs9DvaAkSVCtFFALqwBvnscTNnUXLB3xxX1Is/edit#)). However, when two non-linear causal environments were simulated (Case 2), they were not ranked as most important when all alleles were considered (see Figure **S5**).

GF Offset had a consistent negative correlation with CG Fitness across all replicates for all four multilocus cases, in both Core and Edge demes, with similar performance regardless of whether all loci or only causal loci were used (orange bars, **Fig 4**). While GF Offset was not consistently as well-correlated with CG Fitness as the causal environmental distance (dark green bars, **Fig 4**), it consistently outperformed overall environmental distance (light green bars, **Fig 4**), F_ST Genome_, and F_ST Causal_ (light blue bars, **Fig 4**) as a predictor of CG Fitness.

**Figure 4.**
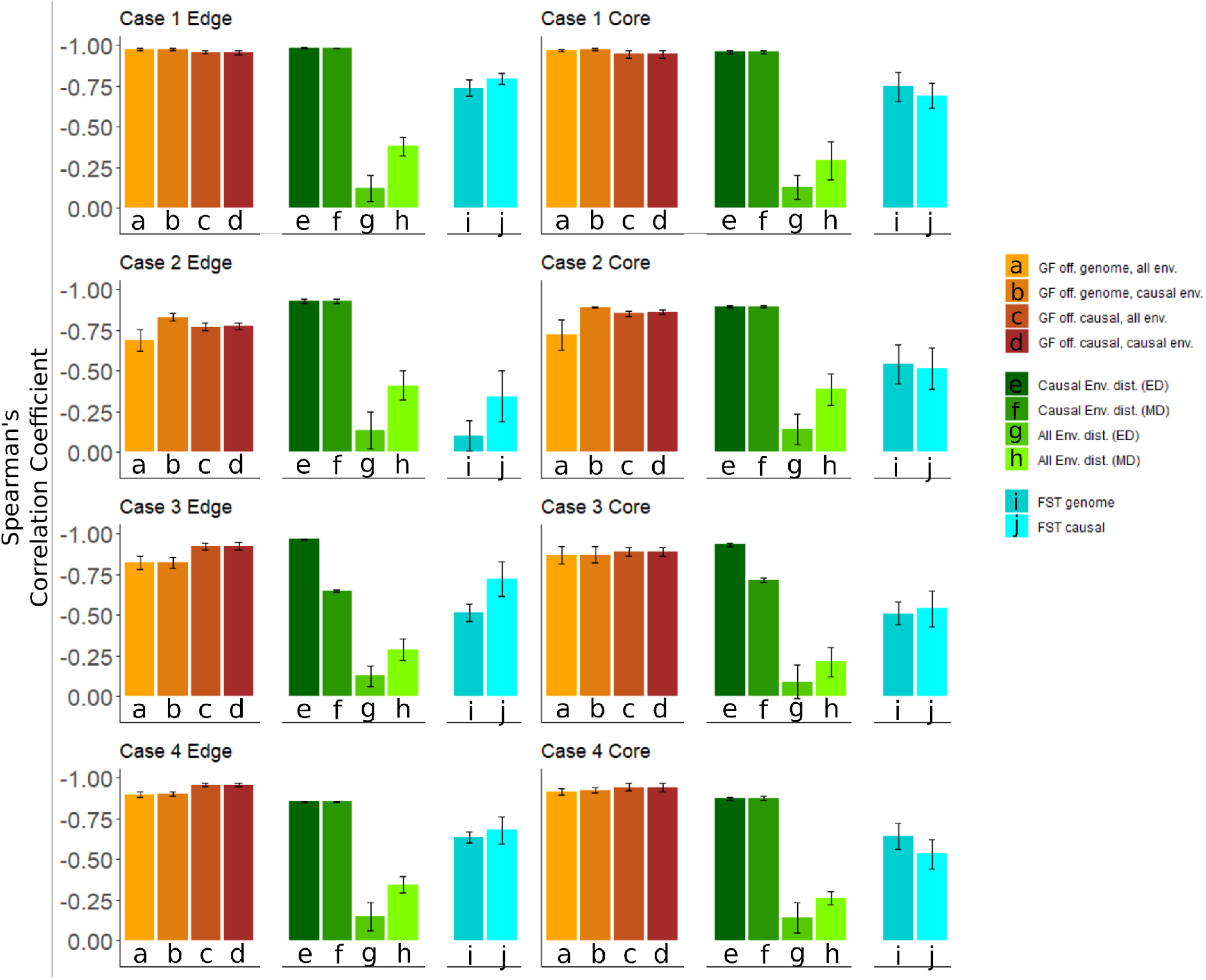
Spearman’s correlation between Common Garden (CG) Fitness and different measures of offset (GF Offset = browns, Environmental distances = greens, *F*_ST_ = blues), for common gardens at the edge of the landscape (left column) and common gardens in the center (core) of the landscape (right column). Each row represents an individual case of the multilocus simulation: first row shows Case 1, where both causal environments were linear orthogonals; second row Case 2, where both causal environments were non-linear orthogonal “mountain peaks”; third row Case 3, where the optimal phenotypic range for trait two was narrower than for trait one; fourth row Case 4, where the strength of selection on trait two was weaker than on trait one.

### GF Offset versus other measures of offset

#### Environmental distance as a predictor of fitness across multilocus Cases

When only the environments underlying local adaptation were considered, *E_D_* and *M_D_* were consistently the most correlated with CG Fitness across all cases of our polygenic model (dark green bars, **Fig 4**). However, when non-causal variables were included in the offset calculations, GF Offset outperformed *E_D_* and *M_D_* by a wide margin. Between the two environmental distance measures, *M_D_* performed just as well as *E_D_* in all cases except Case 3, where the internal standardization of each environment by its variance caused the 2nd environmental variable with lower variance to bias *M_D_*.

#### Genetic distance as a predictor of fitness across multilocus Cases

Measures of *F*_ST Genome_ and *F*_ST Causal_ generally were among the poorest predictors of CG Fitness across all cases, but did perform slightly better than overall environmental distance. Offset predictions by both measures of *F*_ST_ were more impacted by deme location (core vs edge) than any other method. While *F*_ST Causal_ was generally a better predictor of CG Fitness overall for the Edge demes, *F*_ST Genome_ performed similarly in the Core demes. In all simulated multilocus cases, GF was able to accurately determine the underlying causal environments, except when the causal environments were non-linear and non-adapted alleles were included (**Fig S5** - [supp](https://docs.google.com/document/d/1oCI5_FIs9DvaAkSVCtFFALqwBvnscTNnUXLB3xxX1Is/edit#)).

## DISCUSSION

Across all simulated cases GF Offset performed well at predicting CG Fitness, regardless of whether or not non-adapted loci and non-causal environments were included in the analysis. We also found that neutral demography can confound GF Offset, underscoring the need to correct for population structure before training models, and that GF Offset can be sensitive to sampling schemes. GF Offset can be representative of changes in fitness under both the simulated single-locus and polygenic architectures, lending support to the key assumption of a negative relationship between GF Offset and fitness that underlies the use of GF for forecasting maladaptation. When all environmental variables were considered (causal and non-causal), GF Offset, which is based on weighting of environmental gradients given the strength of their association with adaptive variation, outperformed predictions of changes in fitness from the unweighted distance metrics. However, when environmental drivers of adaptation were known and only those gradients were included in the offset calculation, environmental distance performed as well as, or better, than GF Offset.

### Interpretation and comparisons

The strength of the relationship between causal environmental distance and CG Fitness underscores the need to identify environmental drivers of local adaptation. While one study has found a positive relationship between the strength of genotype–environment associations and environments that predict common garden fitness (Mahony *et al*. 2020), a strength of GF is its ability to identify linear selective environments from multiple candidate environmental variables. Although this was not true in Case 2 when all alleles were included (see **Fig S5** - [supp](https://docs.google.com/document/d/1oCI5_FIs9DvaAkSVCtFFALqwBvnscTNnUXLB3xxX1Is/edit#)). GF Offset consistently was a better predictor of CG Fitness than *F*_ST_, regardless of whether *F*_ST_ was calculated using only adapted loci or not. This has significant implications for the use of *F*_ST_ as a decision metric for prioritizing conservation efforts (Xuereb *et al*. 2020). It should be noted that the magnitude of GF Offset can not be compared across different studies, as there is no currently accepted approach to standardize the measure (e.g., to account for differences in the number of variables used in the analysis, etc. But see [supplemental results](https://docs.google.com/document/u/0/d/1oCI5_FIs9DvaAkSVCtFFALqwBvnscTNnUXLB3xxX1Is/edit) in **S1**).

### Conceptual concerns

Although GF Offset is increasingly used to forecast maladaptation, we still have a poor understanding of its performance in natural systems. The underlying assumptions in predictive applications of GF Offset, as with most other approaches to fitness inference, are that a population already occupies its adaptive optimum when sampled and therefore changes in allele frequency composition (regardless of direction) result in decreases in fitness. Furthermore, GF Offset assumes that the molecular signatures of local adaptation when multiple demes occupy the same environment at different sites should be the same. However, this is further predicated on the stability of both the adaptive landscape and the genomic architecture maintaining fitness. These assumptions may be violated, with shifts in the adaptive landscape potentially driven by fluctuations in climate and ecology (Arnold *et al*. 2001), and under transient, highly redundant genomic architectures (Láruson *et al*. 2020). With the single locus of large effect simulation showing the strongest negative correlation between GF Offset and CG Fitness, and the non-linear environment simulation (Case 2) the lowest, GF Offset is clearly impacted by both genomic architecture and the nature of the common garden considered. Also, for linear allele frequency clines, the steepness of the cline did not influence the (non-linear) cumulative importance curve. Although when the cline was non-monotonic, the cumulative importance curve better matched the pattern of turnover. Therefore, at least for linear clines, the interpretation of the slope of the cumulative importance curve as a measure of the rate of allele frequency change may not be appropriate.

### Caveats & best practices

The sensitivity of GF Offset to deme size requires special consideration when studying populations that do not maintain a uniform distribution across their range, especially for populations of conservation concern (Rellstab *et al*. 2015; Borrell *et al*. 2020). Empirical studies have found negative associations between GF Offset and population size (Bay *et al*. 2018; Ruegg *et al*. 2018), but our results show that these associations can arise due to neutral genetic drift and not signals of selection as assumed. At small (large) deme sizes there is more (less) genetic drift, which leads to greater (less) allele frequency turnover at that end of the environmental gradient, and therefore more (less) rapid increases in cumulative importance and larger (smaller) GF Offset. These results highlight that empirical observed negative relationships between GF Offset and population size cannot be assumed to indicate a selection-driven response, and underscores the need to account for population structure when fitting GF. Since other metrics of genetic offset have been found to be associated with population size (Borrell *et al*. 2020), the potential effect of genetic drift on various offset measures should be more fully evaluated. To this end, recent studies have explored correcting allele frequencies for population structure based on the population covariance matrix (Berg & Coop 2014) prior to analysis with GF for outlier detection (Fitzpatrick *et al*. 2021). Empirical studies that find a correlation between population size and offset values should not be considered examples of validation for offset measures. Additionally, datasets where deme size and the most important predictor environment are correlated would be most susceptible to this phenomenon, and investigators should report these relationships.

In our simulations the degree of negative correlation between GF Offset and fitness depends on genetic architecture, the genotype-phenotype-fitness map, and the pattern of environmental variation on the landscape. The thought experiments showed that GF can be sensitive to sampling schemes, and has higher performance when populations are densely sampled along environmental gradients, raising questions about how sampling schemes might bias environmental predictor importance values, which requires further study. Sampling considerations are further impacted by the way GF trains itself on approximately 2/3 of the input number of populations, so GF’s ability to adequately predict the training data can be impacted when only a few populations are analyzed.

### Limitations of in silico experimentation

To assess the predictive potential of GF Offset, our simulations focused on a select few “idealized” scenarios (i.e., high local adaptation, known causal environments, fitness assessment across all common gardens at a fixed time point), with corresponding data which is unlikely to be reflected in empirical work. All calculations using causal alleles were not dependent on those alleles being identified as outliers – it was simply assumed that they were known. In reality, even though many causal alleles showed relatively elevated *F*_ST_ values (**Fig S6** - [supp](https://docs.google.com/document/d/1oCI5_FIs9DvaAkSVCtFFALqwBvnscTNnUXLB3xxX1Is/edit#)) most would be unlikely to be identified through any outlier cut-off approach. A key issue to highlight is that all fitness values simulated here have been calculated as relative fitness, whereas most conservation-minded applications of GF will be concerned with absolute fitness (i.e., population size may be shrinking with increasing genotype–environment mismatches). This distinction between absolute and relative fitness is critical when using models to inform conservation management decisions, since changes in allele frequencies (due to genetic drift or differences in relative fitness), do not necessarily impact demography as absolute fitness does. In fact, allele frequency changes can *only ever* reflect relative fitness (Brady *et al*. 2019). For example, a novel genotype might be increasing in a deme because it has higher relative fitness than another genotype, but both the demes could still be declining in size because both genotypes have low absolute fitness. Therefore, the rationale that allele frequency changes would be useful in making fitness predictions under climate change need closer examination.

### Conclusions

While most emergent complexities involved in applying GF Offset to realistic scenarios are still poorly understood, there is still promise in the application of this method to identifying key environmental drivers of local adaptation and for estimating fitness declines in response to rapid environmental change. Key considerations of demography, genomic architecture, and the nature of environmental gradients have been highlighted here, as in earlier work (Capblancq *et al*. 2020; Fitzpatrick *et al*. 2021; Gougherty *et al*. 2021), as factors that can have significant effects on measures of GF Offset. All future inferences drawn from the potential negative relationship between GF Offset and fitness must take care to address these features of the study system explicitly, and acknowledge the limitations of all inferences if any of these factors are not well understood.

