## Supplement 1 for "Seeing the Forest for the trees: Assessing genetic offset predictions with Gradient Forest"

### Table of contents

Supplemental Methods
- Single-locus single-environment population genetic simulations
- Multilocus two-trait two-environment quantitative genetic model.

Supplemental Results

Figures
- Figure S1
- Figure S2
- Figure S3
- Figure S4
- Figure S5.
- Figure S6

### Supplemental Methods

#### *Single-locus single-environment population genetic simulations*

Fitness was modeled as a linear function of a stable environmental gradient ($E_i$) for the three genotypes:

$$\omega_{AA} = 1 + s * E_i$$

$$\omega_{aa} = 1 - s * E_i$$

$$\omega_{aA} = (\omega_{AA} + \omega_{aa})/2 = 1$$

where *s* is the fitness slope parameter, and values of $E_i$ follow a linear cline across the x-axis of the landscape (from −1 to 1). Note that there is no dominance (the heterozygote is intermediate to the two homozygotes), which resulted in individuals who were heterozygous, possessing one derived and one ancestral allele, being treated as functionally neutral with respect to the environment.

All individuals in the simulation were initialized with the ancestral *a* allele. The derived allele *A* was added at a single point of origin in one copy of one individual's genome, located in $D_{2,9}$ where it increased fitness, and was then allowed to move through the metapopulation for 300 generations. Since all neutral variation was added *post-hoc* via mutation overlay on recapitated coalescent ancestry, as described earlier, no burn-in period was implemented prior to the introduction of the derived allele. The simulation was replicated ten times. At the end of the simulation, we evaluated the ability of GF offset to predict the mean fitness of demes when their individuals were transplanted to new environments, as described in the main text.

#### *Multilocus two-trait two-environment quantitative genetic model.*

In order to let the populations build up some additive genetic variance, the optimum for each phenotype *P* in each deme *j* ($\Theta_{1jt}$ and $\Theta_{2jt}$) was gradually fixed over time (*t*) after a burn-in period. Optima were kept at 0 for the first 1000 generations, and subsequently increased gradually to their final values over another 1000 generations. During the initial burn-in, stabilizing selection was kept weak ($\sigma_k = 4$) for all populations to allow additive genetic variance to build, followed by a gradual narrowing of the multivariate normal distribution to the final value (see section next section).

The homogeneous initial burn-in, for generations t = {1-1000}, for all *j*, was calculated with

$$\Theta_{1jt} = \Theta_{2jt} = 0$$

$$\sigma_k = 4$$

while the transition subsequent burn-in, for generations *t* = {1000–2000}, was calculated with

$$\Theta'_{jt} = \Theta_{jt}\left(\frac{t - 1000}{1000}\right)$$

$$\sigma'_k = \sigma_k * \ 1 - \left(\frac{t - 1000}{1000}\right)$$

After 2000 generations, the optima were set to a constant value for each deme (see next section).

### Supplemental Results

The magnitude of GF Offset across different studies is not readily comparable, as there is no currently accepted approach to standardize the measure (e.g., to account for differences in the number of variables used in the analysis, etc). However, we here list the values measured across our different simulations for the reader to get a general sense of the scale of GF Offset values.

*Neutral sims GF Offset values*: Maximum GF Offset values observed across the different replicates for each scenario were 0.106 when demes were uniform, 0.096 when deme size increased, and 0.117 when deme size decreased.

*Single locus Gf Offset values*: Maximum GF offset values were seen when the causal allele was considered, with little difference between all environments being considered (1.06696) or only the causal environments (1.06686). A similarly small difference was seen when all alleles were considered and all environments (0.069) and only causal environments (0.0686).

*Multilocus GF Offset values*: Case 1 had the highest GF Offset values (0.112), followed by Case 3 (0.0207), Case 4 (0.0198), and Case 2 (0.00467). Maximum GF Offset values were observed when only causal alleles and all environments were considered.

Figures

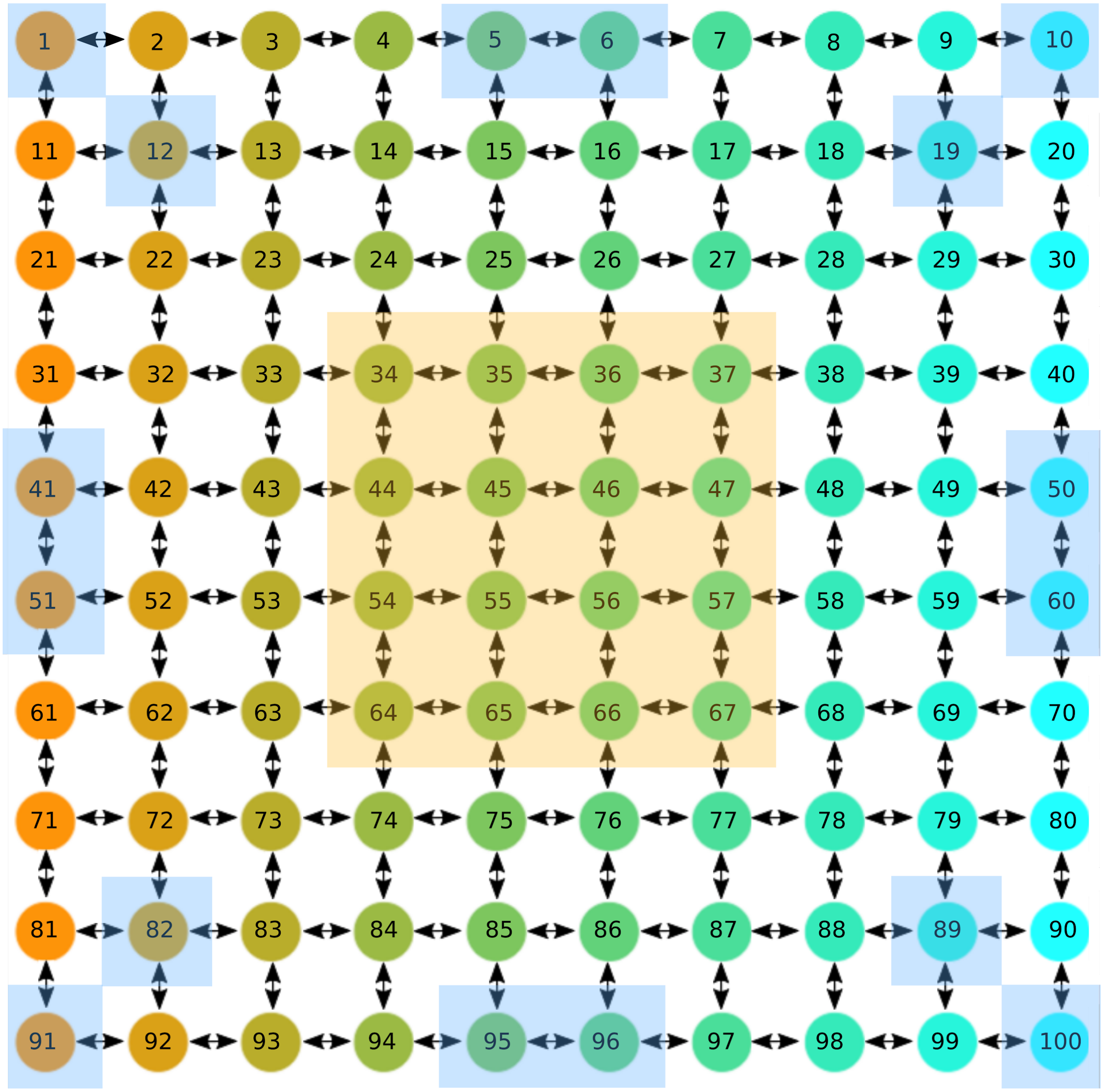


**Figure S1**

Visualization of the 100 demes in the meta-population model. When simulating a longitudinal cline (in the single locus model and the first environment of the multilocus model), each column of demes shared a single environmental value, increasing from the West (orange, column 1, environment -1) to East (cyan, column 10, environment +1). Edge demes used in the multilocus comparison to common garden fitness are highlighted in light blue rectangles, while Core demes are highlighted in orange rectangles.

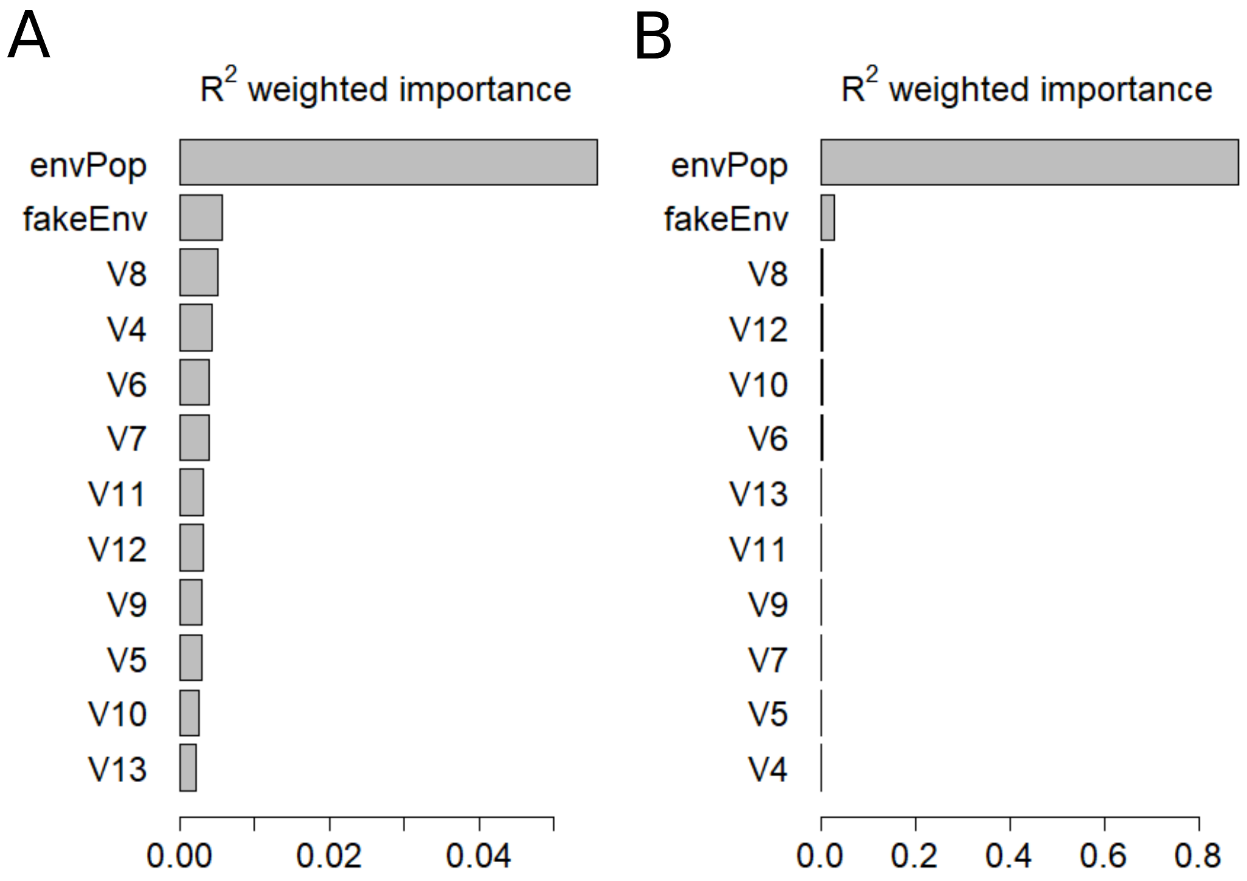


**Figure S2**

GF values of $R^2$ weighted importance for environmental predictor variables in the single locus simulation. (**A**) were calculated with the whole genome datasets, i.e. not filtered for adapted alleles. (**B**) were calculated using only the environmentally adapted allele. Environment "envPop" is the causal environment in the simulation, while "fakeEnv" is a non-causal environment correlated with the causal environment, and "V" desingates random environmental values.

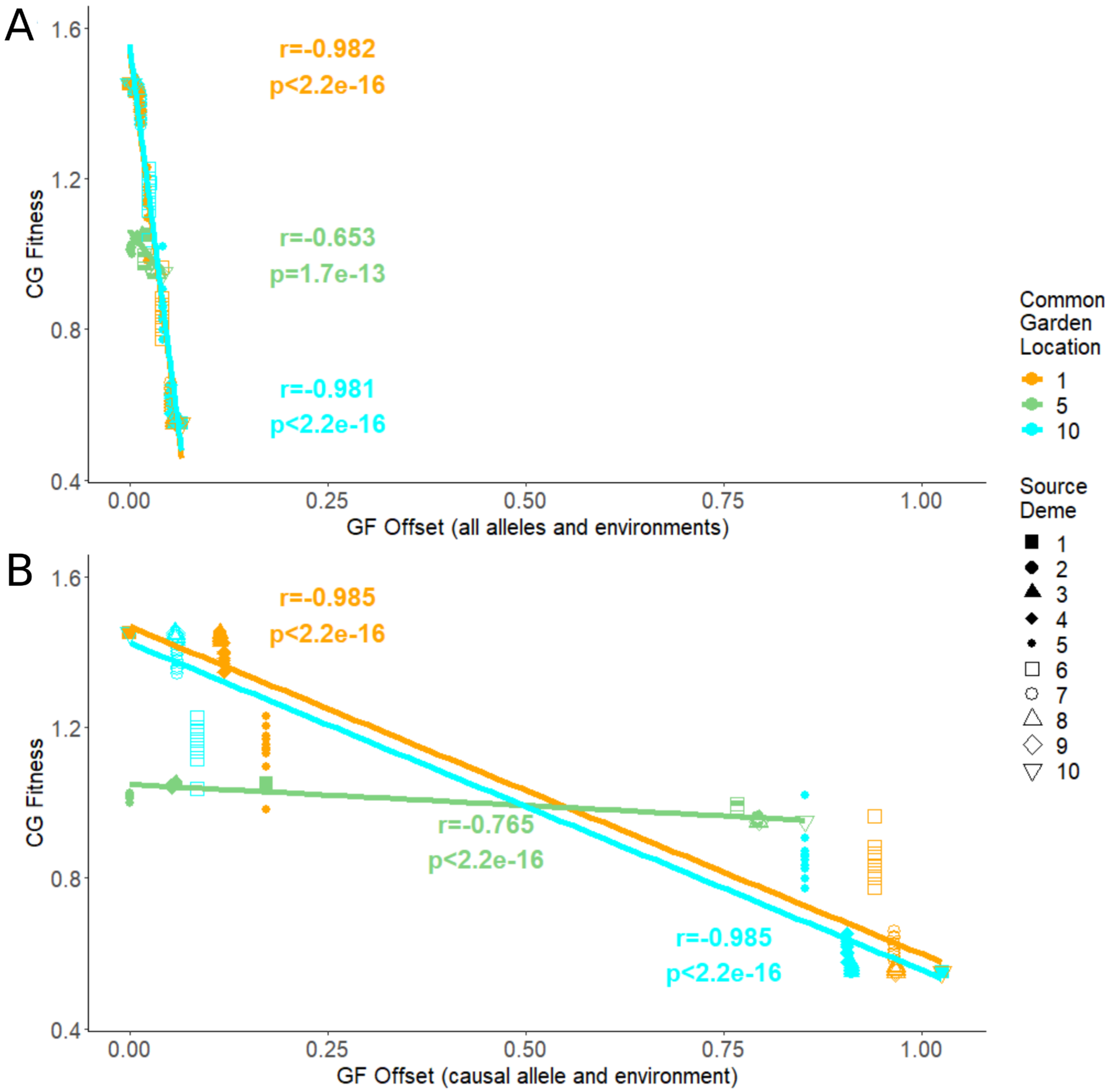


**Figure S3**

GF Offset from the single locus "population genetic" model versus Common Garden fitness, with (A) GF Offset calculated across the whole genome and all environments and (B) with GF Offset calculated only from the single causal allele and environment. In the single locus model, all demes within each column of the metapopulation matrix share the same environment; numbers 1 through 10 in the figure represent the columns in that matrix.

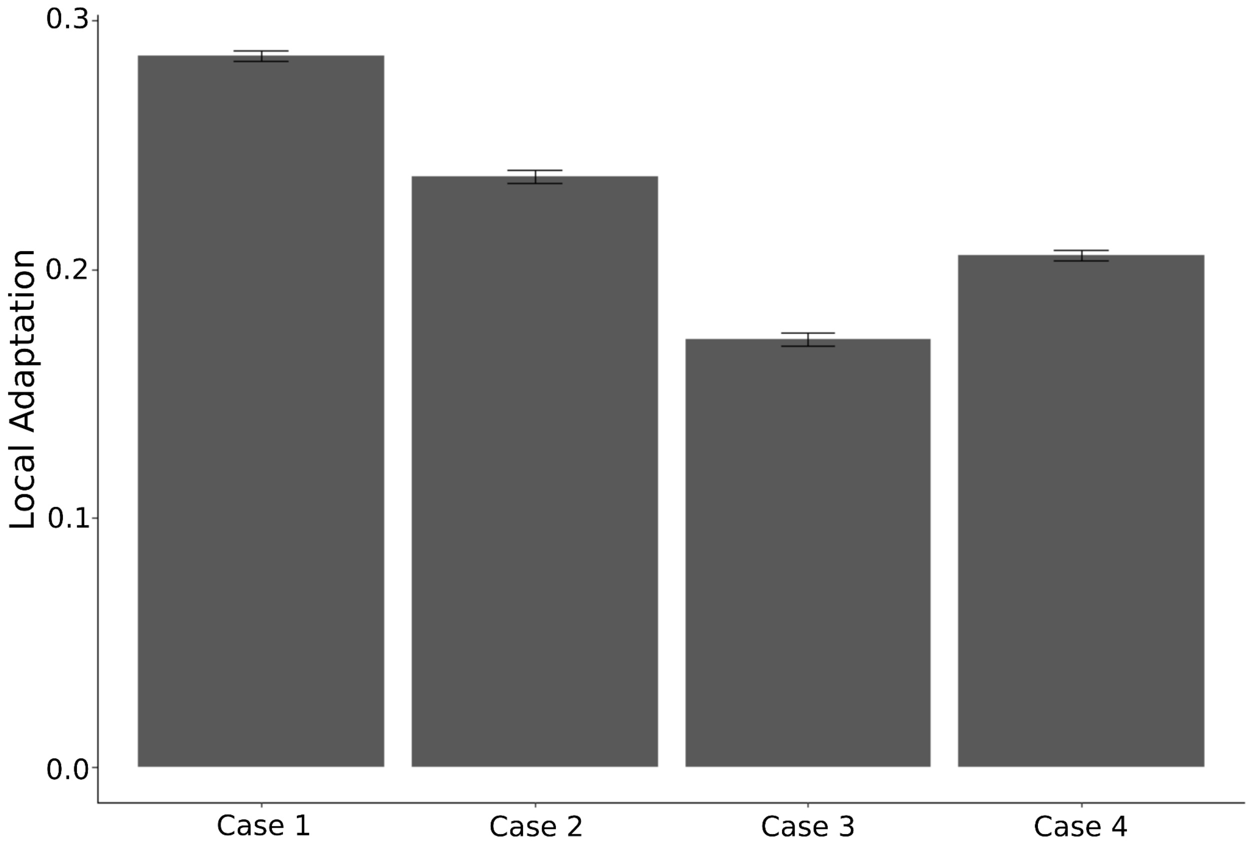


**Figure S4**

Comparison of magnitude of local adaptation attained in each simulated case across the multilocus simulations, with standard deviations across replicates shown. Case 1 (two linear causal environments) had the highest instance of local adaptation, with an individual on average being 28.8% more fit in their home environment than anywhere else, and Case 3 (one of the two causal environments with a much narrower range in optima) having the lowest with an average at home fitness 16.9% greater than anywhere else.

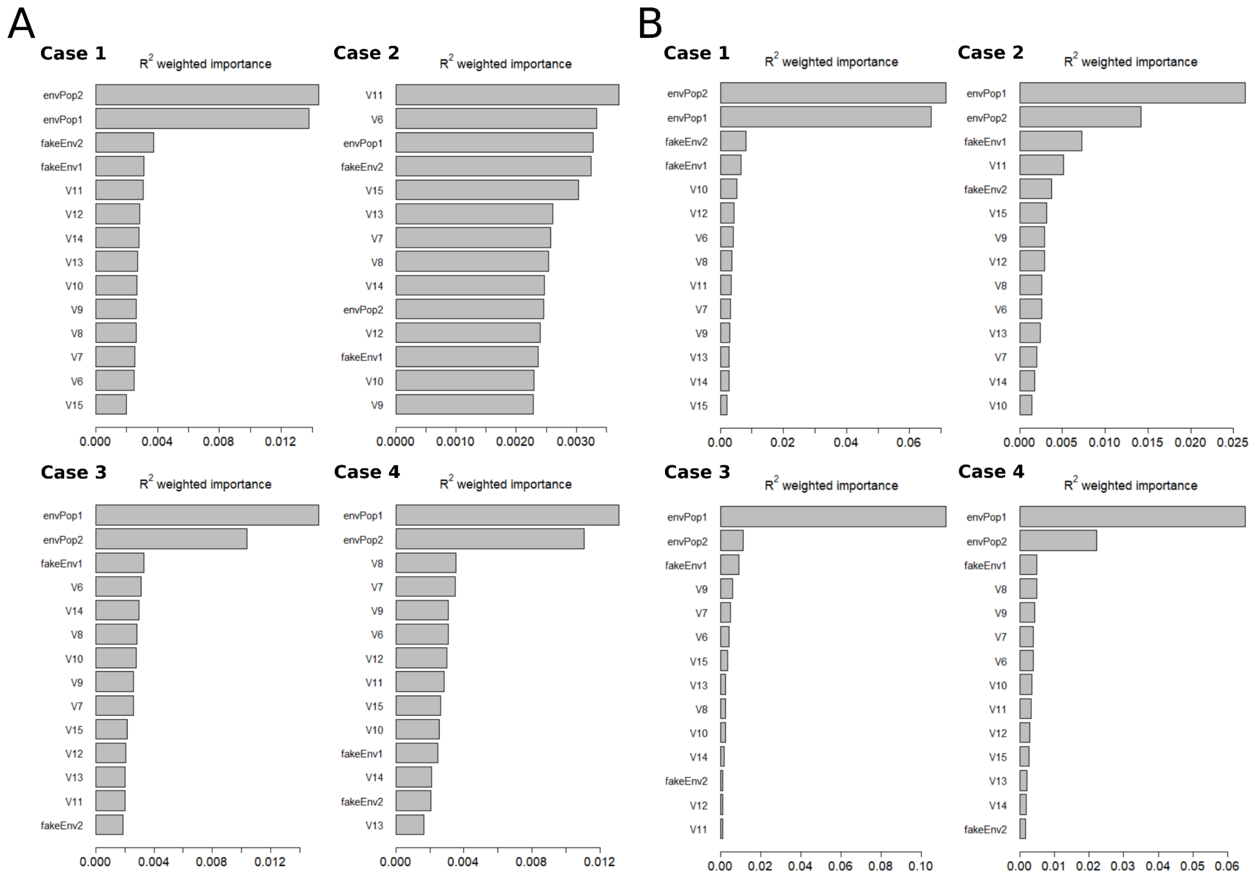


**Figure S5**.

GF values of $R^2$ weighted predictor importance values for one replicate of each of the four multilocus simulation cases. (**A**) were calculated with the whole genome datasets, i.e. not filtered for adapted alleles. (**B**) were calculated with only environmentally adapted alleles. Environments “envPop” are the causal environments in the simulation, while “fakeEnv” are non-causal environments correlated with the designated causal environments, and “V” designates other environmental values. In the non-linear environment Case 2, non-adapted allele $R^2$ signatures confound the individual predictor environment importance values in (**A**). In contrast when only adapted alleles are included (**B**) the individual predictor environment importance values single out the the casual environments.

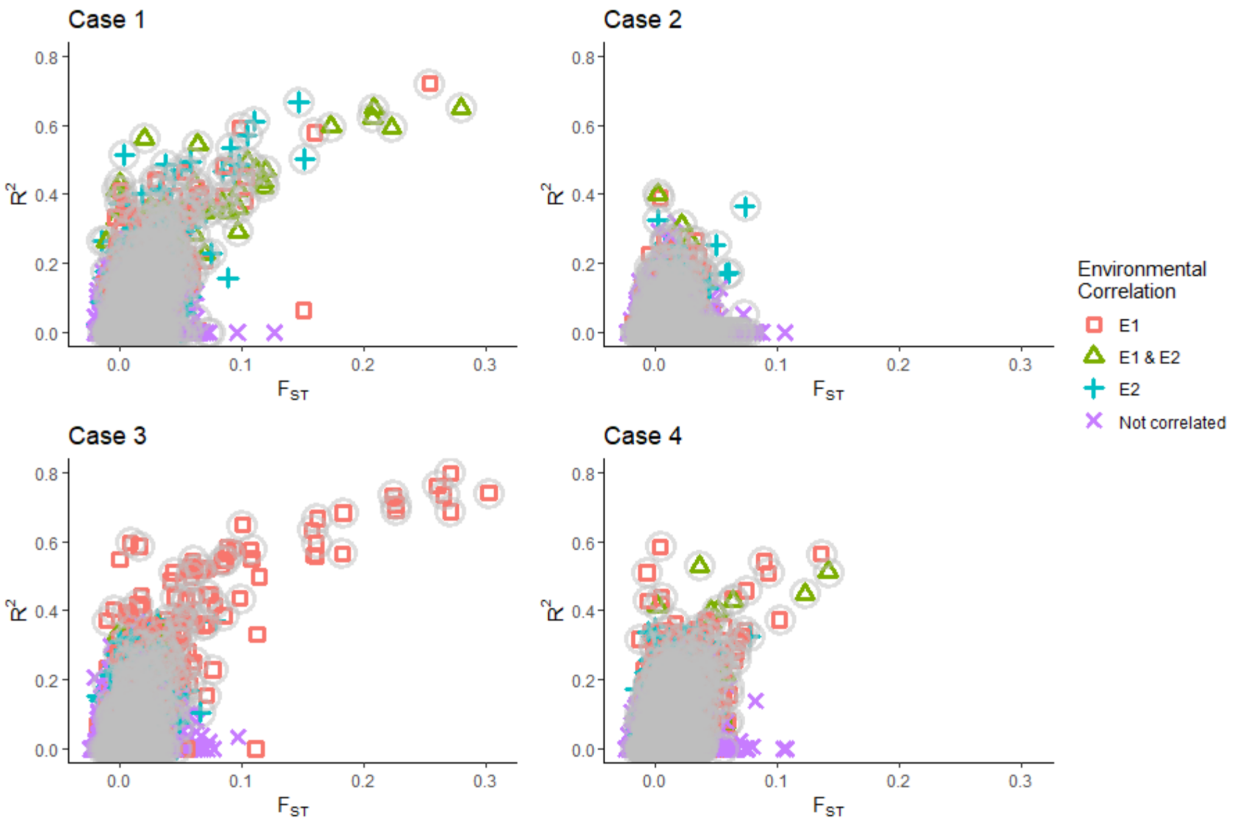


**Figure S6**

Measure of an allele's Pearson's correlation ($R^2$) to an environment in the multilocus simulations, plotted against the per-locus Weir-Cockerham '84 $F_{ST}$ value. Data from all ten replicates were combined for each case. The correlation of each allele to an environment (environment 1, 2, both, or neither) is represented by a unique shape. Alleles under selection are highlighted with a grey circle.
