## Supplement 2 for "Seeing the Forest for the trees: Assessing genetic offset predictions with Gradient Forest"

By KE Lotterhos with input from all authors

5/4/2021

#### R Markdown

This is an R Markdown document. Markdown is a simple formatting syntax for authoring HTML, PDF, and MS Word documents. For more details on using R Markdown see http://rmarkdown.rstudio.com.

When you click the **Knit** button a document will be generated that includes both content as well as the output of any embedded R code chunks within the document. You can embed an R code chunk like this:

```
library(gradientForest)
cols <- viridisLite::turbo(10)
pchs <- c(0, 1, 2, 15, 23)
options(warn = -1)
```

#### Run GF model Function

```
runMod <- function(envPop, alFreq, ntree=500, corr.threshold=0.5){
  # envPop is a dataframe of environments
  # alfreq is the pattern of allele frequency across that environment
  alFreq1 <- data.frame(alFreq)
  #names(alFreq1) <- "alFreq1"
  gfMod <- gradientForest(data=data.frame(envPop, alFreq1),
                          predictor.vars=names(envPop),
                          response.vars=names(alFreq1),
                          corr.threshold=corr.threshold,
                          ntree=ntree,
                          trace=F)

  CI <- cumimp(gfMod, "envPop")
  return(list(CI=CI, IsZero=sum(CI$y)==0, R2=gfMod$result))
}

runMod_multi <- function(envPop, alFreq, ntree=500, corr.threshold=0.5){
  # envPop is a dataframe of environments
  # alfreq is the pattern of allele frequency across that environment
  alFreq1 <- data.frame(alFreq)
  #save the different environment names
```

```
  pred.vars<-names(envPop)
  resp.vars <- names(alFreq1)
  gfMod <- gradientForest(data=data.frame(envPop, alFreq1),
                          predictor.vars=pred.vars,
                          response.vars=resp.vars,
                          corr.threshold=corr.threshold,
                          ntree=ntree,
                          trace=F)
  #create a list of CI for each environment and an IsZero entry for each one as well
  CI_env <- list()
  CI_locus_env <- list()
  IsZero <- list()
  for(i in 1:length(pred.vars)){
    CI_env[[pred.vars[i]]]<-cumimp(gfMod, pred.vars[i], type="Overall")
    IsZero[[pred.vars[i]]]<-sum(CI_env[[pred.vars[i]]]$y)==0
    CI_locus_env[[pred.vars[i]]]<-cumimp(gfMod, pred.vars[i], type="Species")
  }
  #save as a list of lists, including the overall CI (gfMod$Y)
  return(list(CI_env=CI_env, CI=gfMod$Y, IsZero=IsZero, R2=gfMod$result, CI_locus_env))
}
```

Output 1: *Cumulative importance prediction as a function of the environment.* We found that in sparse sampling (see below), this sometimes output all 0's

Output 2: *R2* This is the proportion of variance in the allele frequency that is explained by the model.

### First comparison: few vs. many populations sampled along a cline

#### First take home message: GF outputs depends on sampling scheme

Sometimes in pop gen we only have a few populations sampled. Here we explore the consequences of sampling design on GF outputs. All sampling schemes were based on the same relationship betweeen allele frequency and the environment. We sampled 6, 8, 10, 15, or 50 populations across an environmental gradient.

```
## 'data.frame':    6 obs. of  1 variable:
##  $ envPop: num  -1 -0.6 -0.2 0.2 0.6 1
```

The following plot shows the relationship between allele frequency and the environment is the same for the different sampling schemes:

```
par(mar=c(3,1,3,0.4), mfrow=c(5,1), oma=c(4,4,0,0))
plot(envPop_short$envPop, alFreq_L1_short, xlab="environment", ylab="allele freq.", col=cols[1], pch=pc
abline(h=0.5, col=rgb(0,0,0,0.3))
plot(envPop_med$envPop, alFreq_L1_med, xlab="environment", ylab="allele freq.", col=cols[3], pch=pchs[2
abline(h=0.5, col=rgb(0,0,0,0.3))
plot(envPop_med2$envPop, alFreq_L1_med2, xlab="environment", ylab="allele freq.", col=cols[5], pch=pchs
abline(h=0.5, col=rgb(0,0,0,0.3))
plot(envPop_long$envPop, alFreq_L1_long, xlab="environment", ylab="allele freq.", col=cols[7], pch=pchs
abline(h=0.5, col=rgb(0,0,0,0.3))
plot(envPop_long2$envPop, alFreq_L1_long2, xlab="environment", ylab="allele freq.", col=cols[9], pch=pc
abline(h=0.5, col=rgb(0,0,0,0.3))
```

```
mtext("environment", side=1, outer=TRUE, line=2)
mtext("allele frequency", side=2, outer=TRUE, line=1)
```

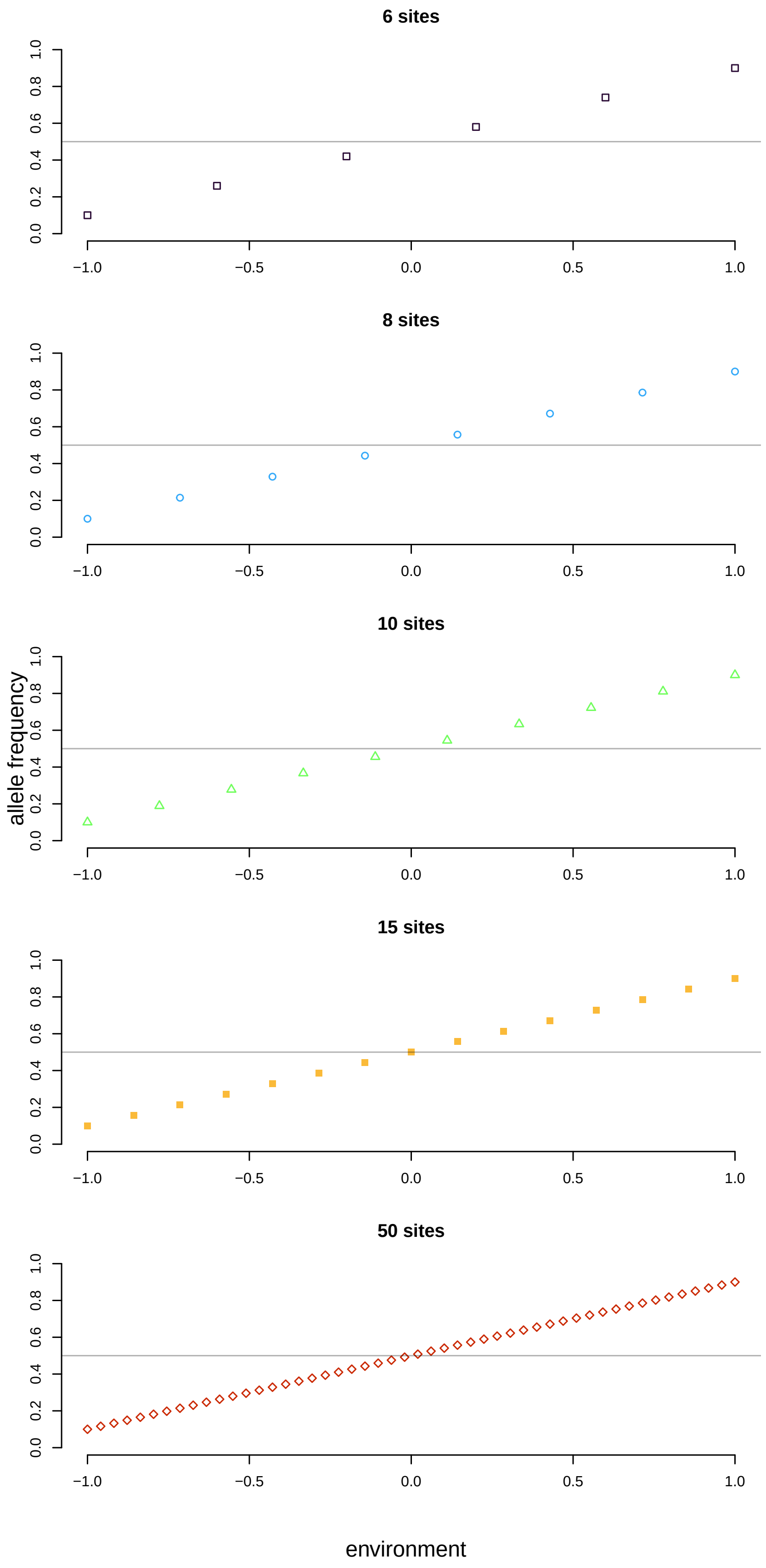
6 sites
8 sites
10 sites
15 sites
50 sites
allele frequency
environment
0.0
0.2
0.4
0.6
0.8
1.0
−1.0
−0.5
0.0
0.5
1.0

For each sampling scheme, we conducted 100 replicate simulations of GF with 500 trees.

The R^2 value from GF is typically interpreted to be similar to a regression. However, it increases with the number of sites sampled along the gradient, as shown in the next graph:

```
# Histogram of R2 values
hist(CI_short_R2, xlim=c(0,1), ylim=c(0,40), col=cols[1], xlab="R^2", main="Distribution of R^2 for\ndi
hist(CI_med_R2, xlim=c(0,1), col=cols[3], add=TRUE )
hist(CI_med2_R2, xlim=c(0,1), col=cols[5], add=TRUE)
hist(CI_long_R2, xlim=c(0,1), col=cols[7], add=TRUE , border = cols[4])
hist(CI_long2_R2, xlim=c(0,1), col=cols[9], add=TRUE )

legend(0, 40, fill=cols[seq(1,9, by=2)], legend = samp_sizes)
```

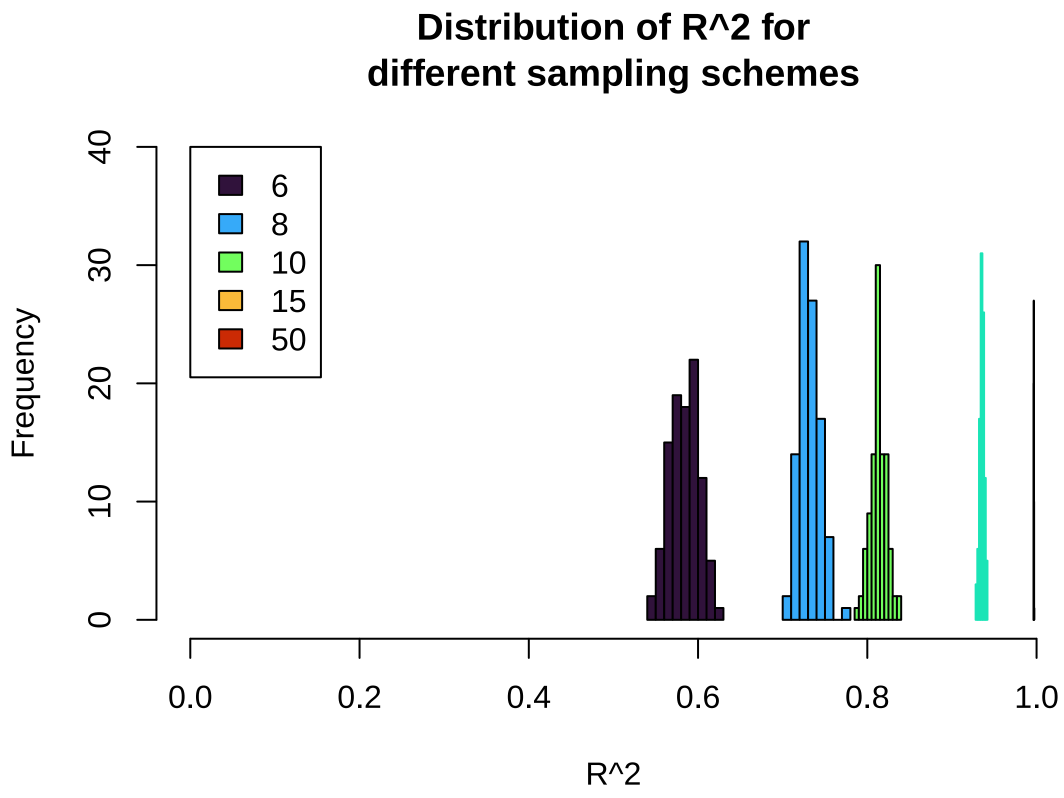


The above two graphs illustrate that the R^2 value from a gradient forests output is not similar to a R^2 value from a regression (which should be the same in all 5 of these cases).

The next figure illustrates that the shape and height of the cumulative importance curve is also sensitive to the sampling design:

```
# CI vs. env
sequ <- seq(1, length(CI_short), by=3)
plot(CI_short[[1]]$x, CI_short[[1]]$y, type="l", col=cols[1], ylim=c(-0.1,1), ylab="CI", xlab="environm
for (i in sequ){
  points(CI_short[[i]]$x, CI_short[[i]]$y, type="l", col=adjustcolor(cols[1], 0.5))
  points(CI_med[[i]]$x, CI_med[[i]]$y, type="l", col=adjustcolor(cols[3], 0.5))
  points(CI_med2[[i]]$x, CI_med2[[i]]$y, type="l", col=adjustcolor(cols[5], 0.5))
```

```
  points(CI_long[[i]]$x, CI_long[[i]]$y, type="l", col=adjustcolor(cols[7], 0.5))
  points(CI_long2[[i]]$x, CI_long2[[i]]$y, type="l", col=adjustcolor(cols[9], 0.5))
}

legend(-0.8, 1, fill=rev(cols[seq(1,9, by=2)]), legend = rev(samp_sizes))
```

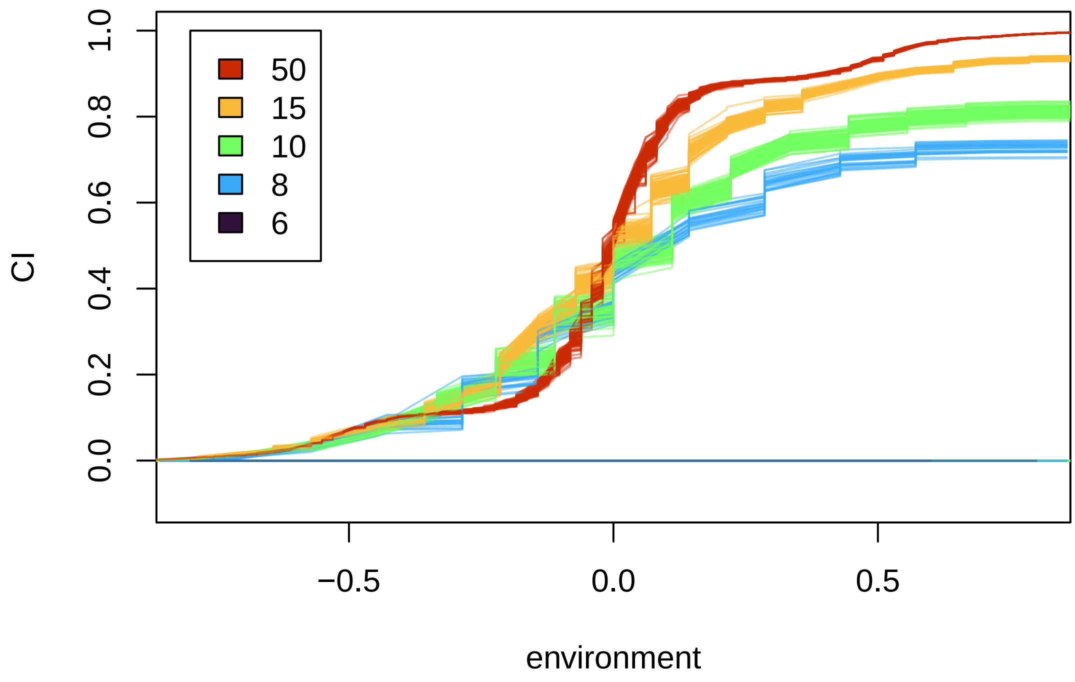


The above plot highlights some interesting behavior.

First, the height of the CI curve depends on the sampling scheme. Sampling more sites along the gradient results in a higher overall CI.

Second, the shape of the CI curve depends on the sampling scheme. Sampling more sites along the gradient results in more sigmodal shaped CI function near the intermediate environment. For example, the actual allele frequency turnover is the same across all 5 sampling schemes. But the CI function makes it look like the sampling scheme with 50 samples is very steep near `environment=0` compared to the other schemes. This highlights issues with interpreting GF outputs as allele frequency change along a gradient and calculating GF offsets from that.

Third, there is an behavior with the distribution of CI values, but it's hard to see. With the sparsest sampling scheme (6 sites), the CI is always 0. With less sparse sampling schemes (e.g., 8 or 10 sites), the CI is *sometimes* 0, but it's hard to see because all the lines overlap at the horizontal line at y=0. We will visualized that in the next plot:

```
# Proportion of 0s
par(mar=c(4,4,3,0.4), mfrow=c(1,1), oma=c(0,0,0,0), bty="l")
plot(samp_sizes, c(CI_short_prop0, CI_med_prop0, CI_med2_prop0, CI_long_prop0, CI_long2_prop0), xlab="nu
```

```
    type="l", col="blue", ylab= "proportion",
    main="proportion of times the CI gives all 0s", ylim=c(0,1))
```

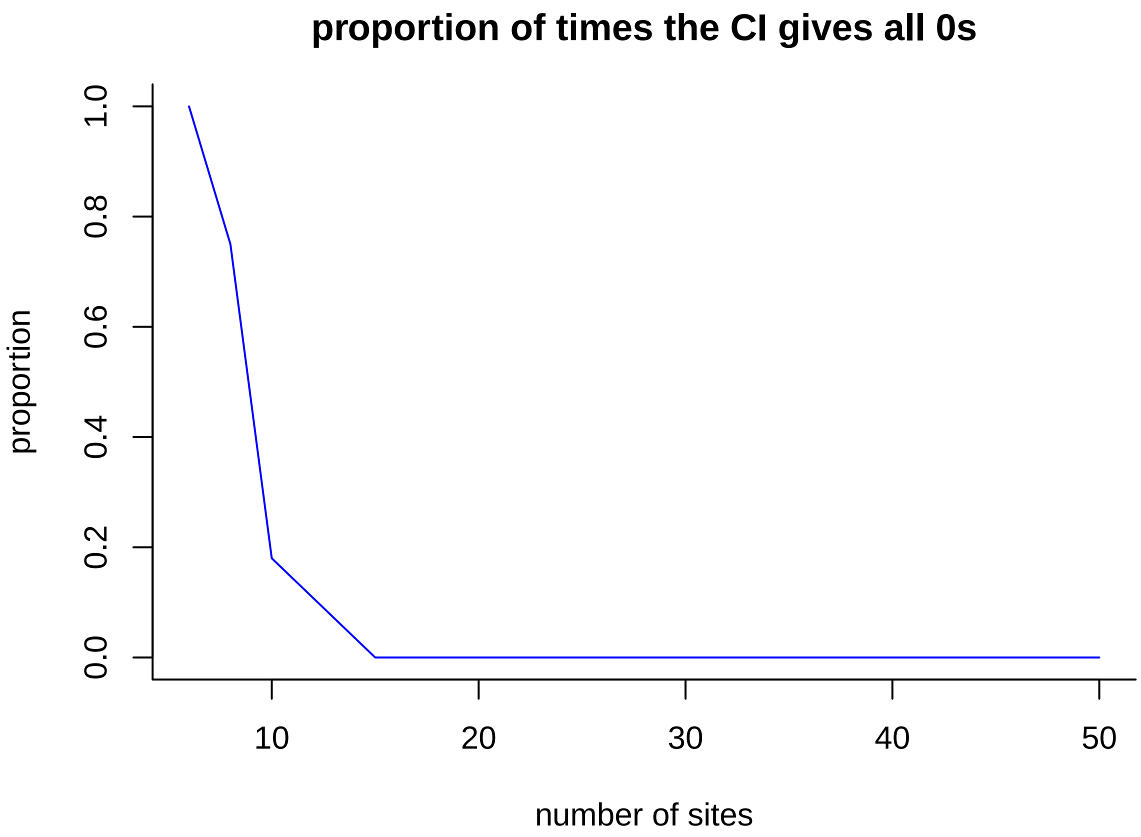


The above plot shows that sometimes GF output all 0s for the cumulative importance function, and that the proportion of times this happened was higher for sparser sampling schemes. We did not find increasing the number of trees changed these proportions.

Another way to visualize it as the distribution of R^2 vs. maximum CI for each replicate simulation:

```
sequ <- seq(1, length(CI_short), by=3)
plot(CI_short[[3]], max(CI_short[[1]]$y), col=adjustcolor(cols[1], 0.5), ylim=c(-0.1,1), xlab="R2", yla
for (i in sequ){
  points(CI_short[[i+2]], max(CI_short[[i]]$y),  col=adjustcolor(cols[1], 0.5), pch=pchs[1])
  points(CI_med[[i+2]], max(CI_med[[i]]$y),  col=adjustcolor(cols[3], 0.5), pch=pchs[2])
  points(CI_med2[[i+2]], max(CI_med2[[i]]$y),  col=adjustcolor(cols[5], 0.5), pch=pchs[3])
  points(CI_long[[i+2]], max(CI_long[[i]]$y), col=adjustcolor(cols[7], 0.5), pch=pchs[4])
  points(CI_long2[[i+2]],max(CI_long2[[i]]$y),  col=adjustcolor(cols[9], 0.5), pch=pchs[5])
}

legend(0.1, 1, col =rev(cols[seq(1,9, by=2)]), legend = rev(samp_sizes), pch=rev(pchs))
```

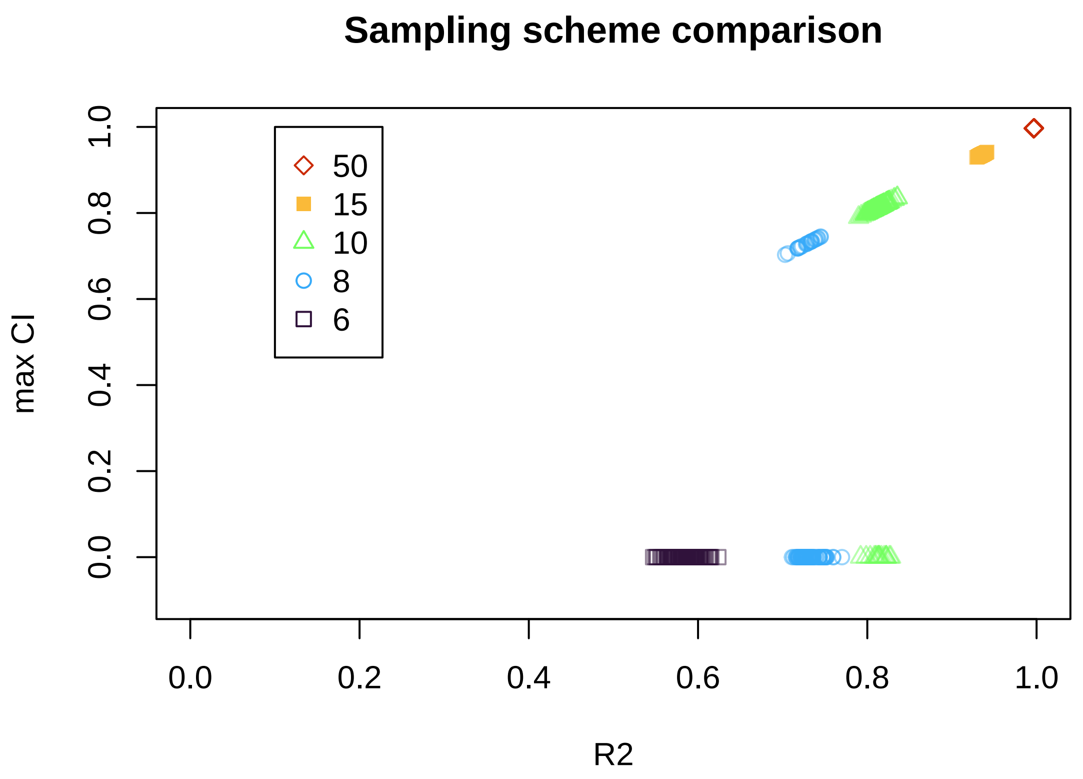


Maybe this weird behavior is GF's way of weighting individual loci that it has more confidence in (e.g, denser sampling). However, it does raise issues for the way that populations are sampled unequally along gradients in nature. In our simulations we had equal number of sites sampled along each environmental gradient (e.g., a 10 x 10 matrix with latitudinal gradient and a longitudinal gradient), so the effect of unequal sampling along different gradients warrants further investigation.

#### Second comparison: A steep cline vs. less steep cline vs. a reverse cline

##### Second take home message: GF outputs are the same regardless of slope of cline

For this comparison, we assumed equal sampling (15 sites along the cline) and manipulated the pattern of allele frequency along the cline.

Code used from first section:

```
envPop_long <- data.frame(envPop=seq(-1,1,length.out=15)) #create env
#create four allele frequency gradients of varying steepness
alFreq_L1_long<-seq(0.1,0.9,length.out=15)
CI_long <- replicate(100,runMod(envPop_long, alFreq_L1_long))
(CI_long_R2 <- (unlist(CI_long[seq(3,length(CI_long), by=3)])))
```

```
alFreq_L1_rev<- rev(alFreq_L1_long)
CI_rev <- replicate(100,runMod(envPop_long, alFreq_L1_rev))
```

```
CI_rev_R2 <- (unlist(CI_rev[seq(3,length(CI_rev), by=3)]))

alFreq_L1_shallow<-seq(0.4,0.6,length.out=15)
CI_shallow <- replicate(100,runMod(envPop_long, alFreq_L1_shallow))
CI_shallow_R2 <- (unlist(CI_shallow[seq(3,length(CI_shallow), by=3)]))
```

As you will see below, all of these clines have the same R^2 value and same max CI (e.g., the same proportion of variance explained by the model). The results highlight, however, why the GF output cannot be called a "vulnerability": the output is agnostic to the direction of allele frequency change across the gradient.

```
par(mar=c(3,1,3,0.4), mfrow=c(3,1), oma=c(4,4,0,0))
plot(envPop_long$envPop, alFreq_L1_long, xlab="environment", ylab="allele freq.", col=cols[7], pch=pchs
abline(h=0.5, col=rgb(0,0,0,0.1))
plot(envPop_long$envPop, alFreq_L1_rev, xlab="environment", ylab="allele freq.", col=cols[2], pch=pchs[
abline(h=0.5, col=rgb(0,0,0,0.1))
plot(envPop_long$envPop, alFreq_L1_shallow, xlab="environment", ylab="allele freq.", col=cols[10], pch=
abline(h=0.5, col=rgb(0,0,0,0.1))
```

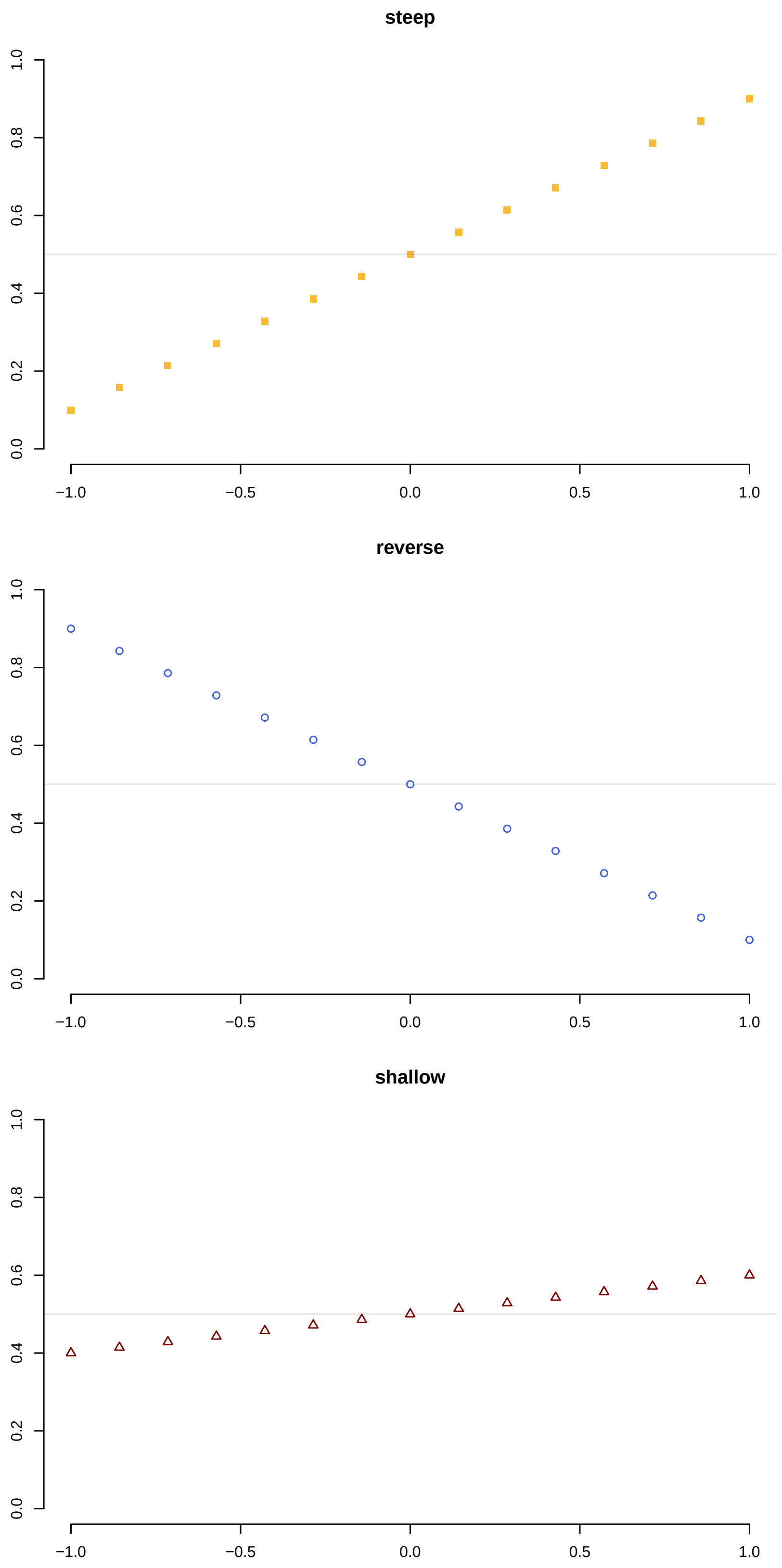
steep
reverse
shallow
0.0
0.2
0.4
0.6
0.8
1.0
−1.0
−0.5
0.0
0.5
1.0

All these relationships give essentially the same CI curve:

```
par(mfrow=c(1,1), mar=c(4,4,1,1))
# CI vs. env
sequ <- seq(1, length(CI_short), by=3)
plot(CI_long[[1]]$x, CI_long[[1]]$y, type="l", col=cols[7], ylim=c(-0.1,1), ylab="CI", xlab="environmen
for (i in sequ){
  points(CI_long[[i]]$x, CI_long[[i]]$y, type="l", col=adjustcolor(cols[7], 0.5))
  points(CI_rev[[i]]$x, CI_rev[[i]]$y, type="l", col=adjustcolor(cols[2], 0.5))
  points(CI_shallow[[i]]$x, CI_shallow[[i]]$y, type="l", col=adjustcolor(cols[10], 0.5))
}

legend(-0.8, 1, fill=cols[c(7,2,10)], legend = c("steep", "reverse", "shallow"))
```

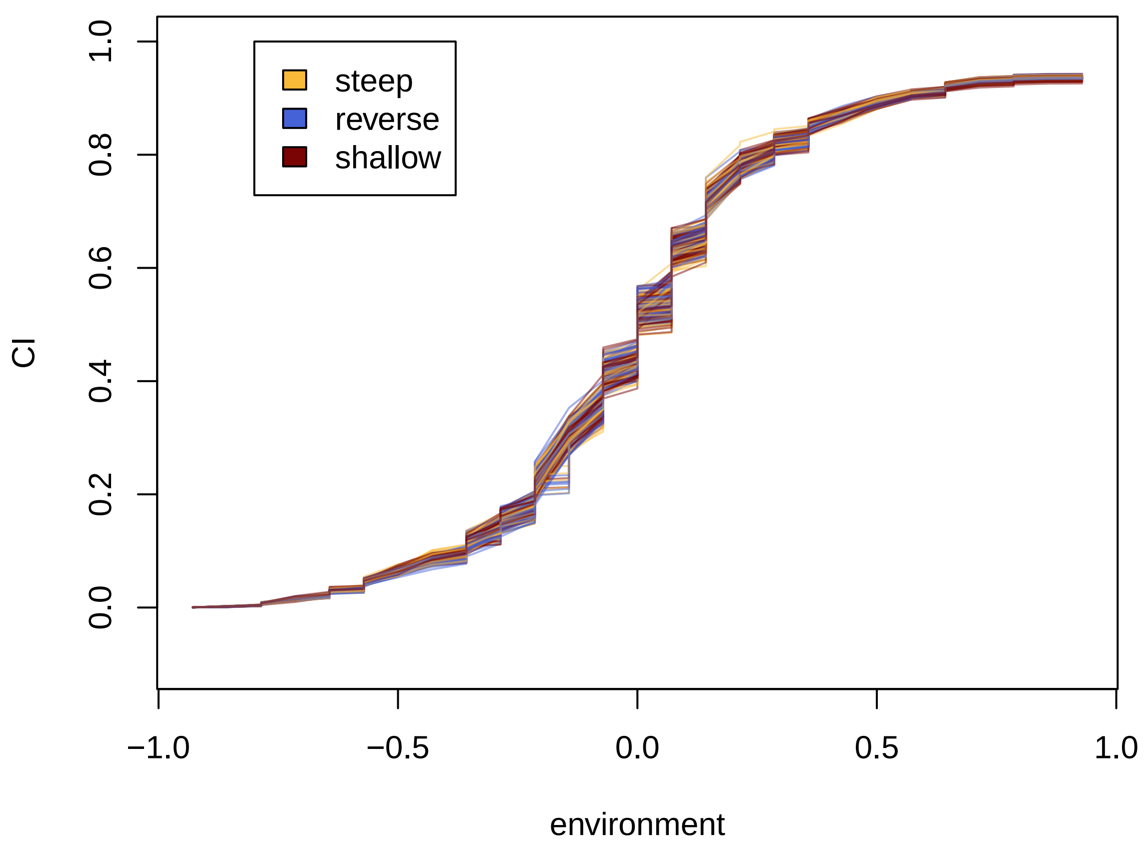


And same R^2 function:

```
hist(CI_long_R2, xlim=c(0.9,1), col=adjustcolor(cols[7], 0.3), ylim=c(0,50), main="Cline type", xlab="R
hist(CI_rev_R2, xlim=c(0,1), col=adjustcolor(cols[2], 0.3), add=TRUE )
hist(CI_shallow_R2, xlim=c(0,1), col=adjustcolor(cols[10], 0.3), add=TRUE )
legend(0.9, 50, fill=cols[c(7,2,10)], legend = c("steep", "reverse", "shallow"))
```

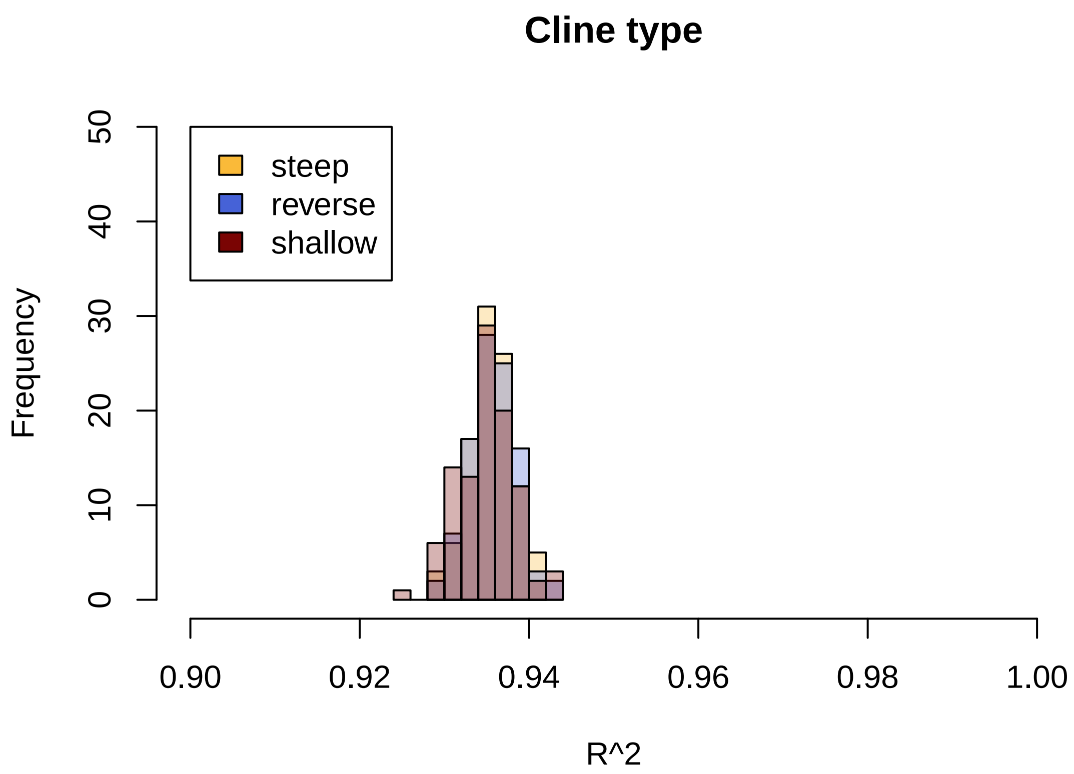


This section highlighted out the GF output is driven by the proportion of variance the model can explain in the data, which is the same for all shapes of allele frequency curves. To further illustrate this, we will consider in the next section changing the shape of the relationship between a.f. and environment.

#### Third comparison: Non-monotonic relationships with allele frequency

##### Third take home message: GF outputs are the same regardless of the magnitude of the non-monotonic relationship with allele frequency, when they have the same shape

For this comparison, we assumed equal sampling (15 sites along the cline) and manipulated the pattern of allele frequency along the cline.

```
envPop_long <- data.frame(envPop=seq(-1,1,length.out=15)) #create env
alFreq_nonMon<- c(seq(0.1,0.9, length.out=5), rep(0.9,5), seq(0.9,0.1, length.out=5))
CI_nonMon <- replicate(100,runMod(envPop_long, alFreq_nonMon))
CI_nonMon_R2 <- (unlist(CI_nonMon[seq(3,length(CI_nonMon), by=3)]))

alFreq_nonMon_rev<- 1-alFreq_nonMon
CI_nonMon_rev <- replicate(100,runMod(envPop_long, alFreq_nonMon_rev))
CI_nonMon_rev_R2 <- (unlist(CI_nonMon_rev[seq(3,length(CI_nonMon_rev), by=3)]))

alFreq_nonMon_shallow<- c(seq(0.4,0.6, length.out=5), rep(0.6,5), seq(0.6,0.4, length.out=5))
```

```
CI_nonMon_shallow <- replicate(100,runMod(envPop_long, alFreq_nonMon_shallow))
CI_nonMon_shallow_R2 <- (unlist(CI_nonMon_shallow[seq(3,length(CI_nonMon_shallow), by=3)]))
```

The shallow non-monotonic case gives the following error in GF, but still runs and produces outputs:

```
Warning in randomForest.default(m, y, ...): The response has five or fewer
unique values. Are you sure you want to do regression?
```

Plot the patterns of allele frequency as a function of the environment:

```
par(mar=c(3,1,3,0.4), mfrow=c(3,1), oma=c(4,4,0,0))
plot(envPop_long$envPop, alFreq_nonMon, xlab="environment", ylab="allele freq.", col=cols[1], pch=pchs[
abline(h=0.5, col=rgb(0,0,0,0.1))
plot(envPop_long$envPop, alFreq_nonMon_rev, xlab="environment", ylab="allele freq.", col=cols[5], pch=p
abline(h=0.5, col=rgb(0,0,0,0.1))
plot(envPop_long$envPop, alFreq_nonMon_shallow, xlab="environment", ylab="allele freq.", col=cols[3], p
abline(h=0.5, col=rgb(0,0,0,0.1))
```

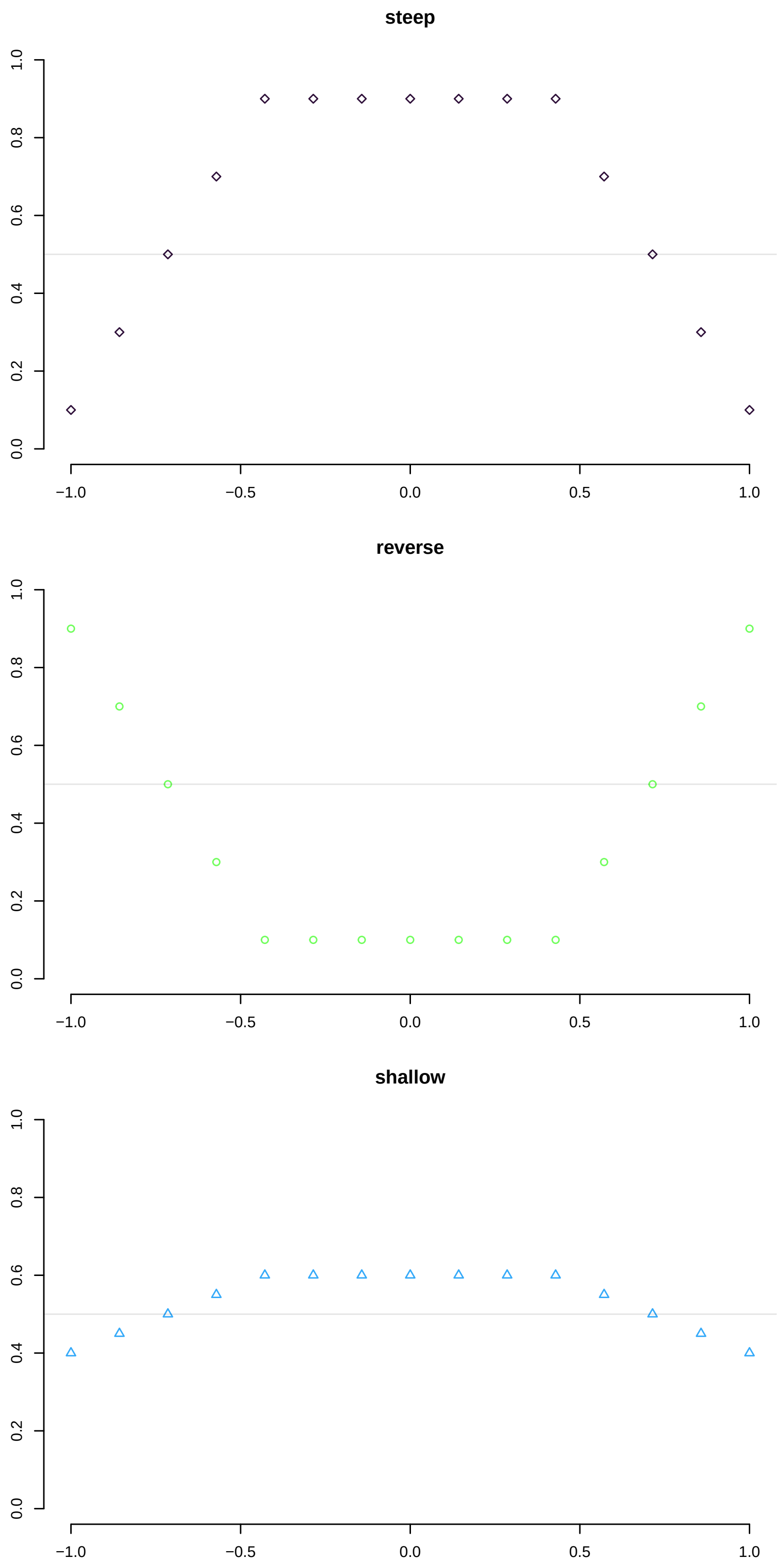

steep
reverse
shallow
0.0
0.2
0.4
0.6
0.8
1.0
−1.0
−0.5
0.0
0.5
1.0

All these relationships give essentially the same CI curve:

```
par(mfrow=c(1,1), mar=c(4,4,1,1))
# CI vs. env
sequ <- seq(1, length(CI_nonMon), by=3)
plot(CI_nonMon[[1]]$x, CI_nonMon[[1]]$y, type="l", col=cols[7], ylim=c(-0.1,1), ylab="CI", xlab="enviro
for (i in sequ){
  points(CI_nonMon[[i]]$x, CI_nonMon[[i]]$y, type="l", col=adjustcolor(cols[1], 0.5))
  points(CI_nonMon_rev[[i]]$x, CI_nonMon_rev[[i]]$y, type="l", col=adjustcolor(cols[5], 0.5))
  points(CI_nonMon_shallow[[i]]$x, CI_nonMon_shallow[[i]]$y, type="l", col=adjustcolor(cols[3], 0.5))
}

legend(-0.8, 1, fill=cols[c(1,5,3)], legend = c("steep", "reverse", "shallow"))
```

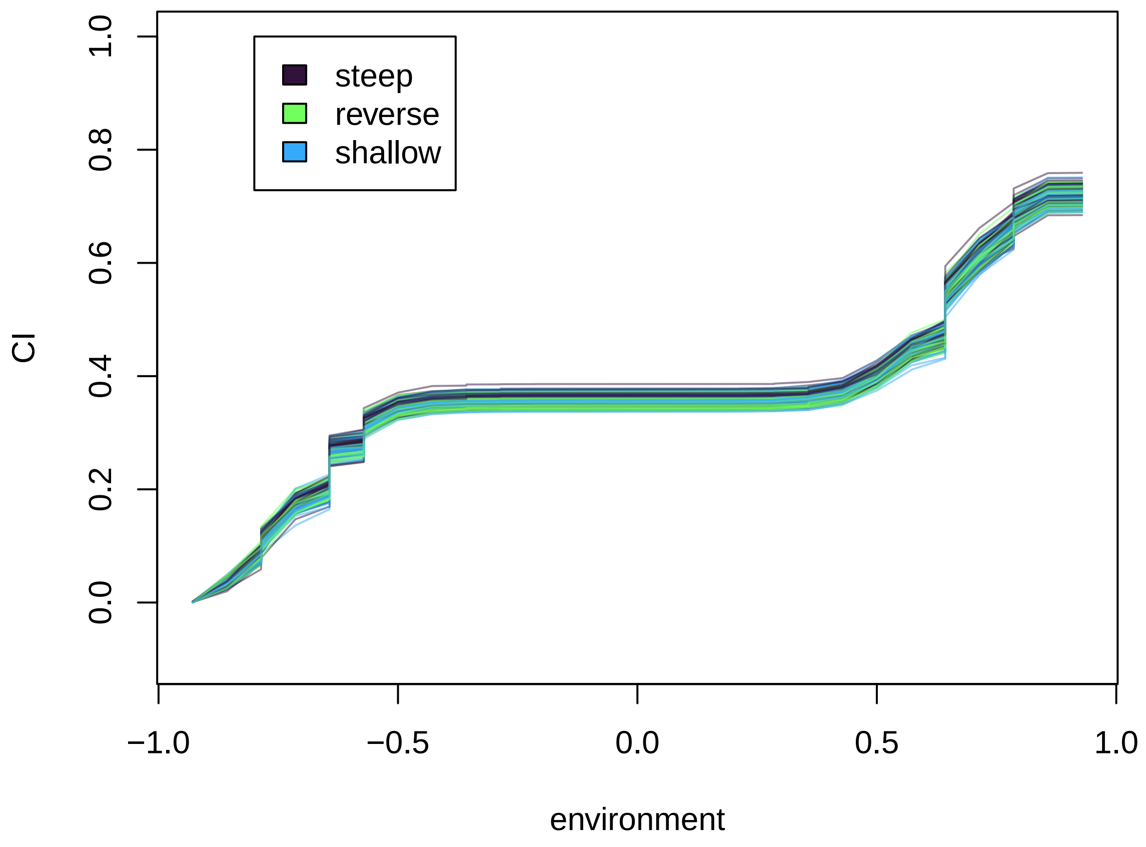


And same R^2 function:

```
hist(CI_nonMon_R2, xlim=c(0.5,1), col=adjustcolor(cols[1], 0.5), ylim=c(0,50), xlab="R^2", main=
      "Distribution of R^2 from\nreplicate non-monotonic sims")
hist(CI_nonMon_rev_R2, xlim=c(0,1), col=adjustcolor(cols[5], 0.3), add=TRUE )
hist(CI_nonMon_shallow_R2, xlim=c(0,1), col=adjustcolor(cols[3], 0.3), add=TRUE )
legend(0.8, 40, fill=cols[c(1,5,3)], legend = c("steep", "reverse", "shallow"))
```

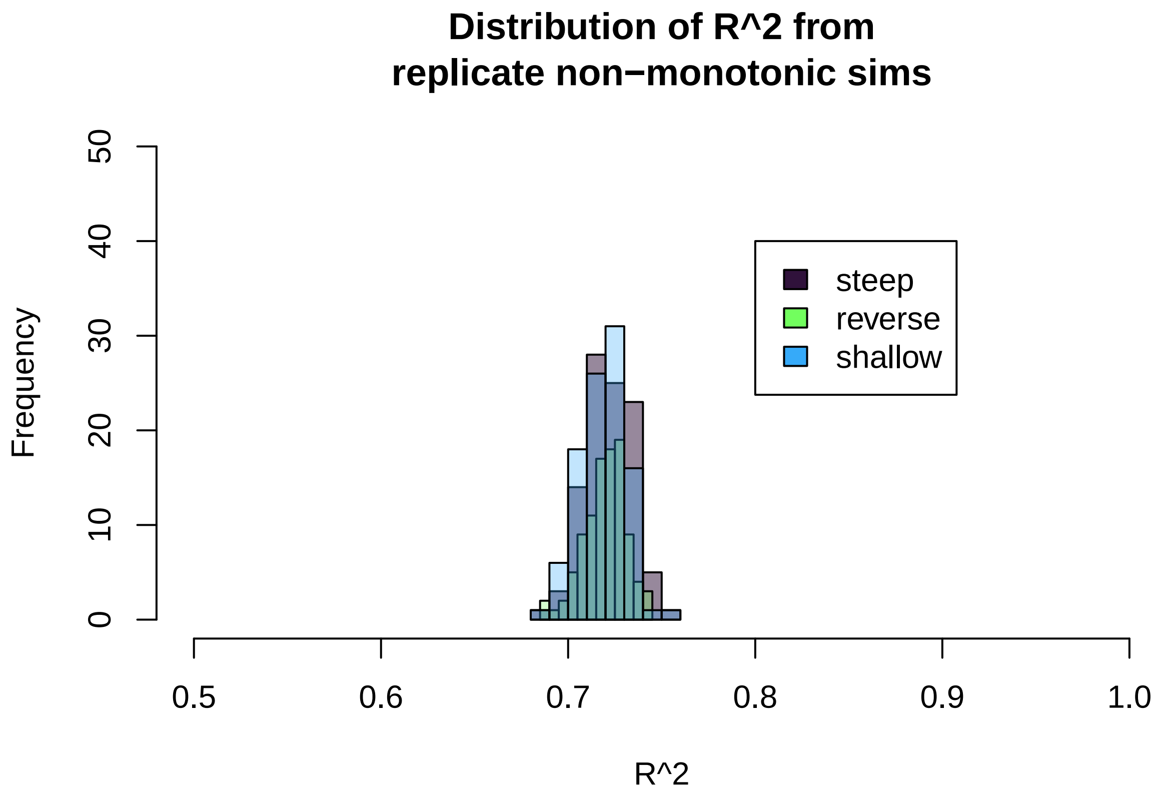


The above two sections illustrate that the GF CI curves do not correspond the amount of turnover in allele frequency. If we consider turnover in allele frequency in the mathematical sense, it is the amount of change in allele frequency per unit change in the environment. In the examples in the above two sections, steeper clines have greater amount of change in allele frequency per unit change in the environment than shallower clines. Hence, if the maximum CI of an allele reflected the turnover, alleles with steeper clines should have greater maximum CI. However, the above two sections show this is not the case when comparing alleles with similar patterns but different amounts of frequency change across the gradient. The maximum CI is, instead, driven by how much variation in allele frequency the gradient-forests model explains, which is the same for steep, shallow, and reverse clines of the same general pattern across the environmental gradient.

The non-monotonic clines in the above example all have the same allele frequency at the extremes of the environmental variable. If the genetic basis of adaptation was monogenic (e.g. caused by a single allele), and fitness followed the same non-monotonic pattern (e.g., individuals with the focal allele had highest fitness in the intermediate values of the environment, and individuals with the alternate allele had highest fitness in the extreme values of the environment), then this conceptual example illustrates issues with calling the GF-offset a genetic offset or genomic vulnerability. In this hypothetical case, individuals of the same genotype would have the same fitness at extreme values of the environment, and hence the genetic offset should be 0. But GF would predict a maximum offset (e.g. the vertical distance in the CI evaluated an environment of -1 and the CI evaluated an environment of +1 in the previous plot).

#### Fourth comparison: A steep cline vs. a non-monotonic cline

##### Fourth take home message: The shape of the CI function reflects the pattern of turnover in allele frequency

For this comparison, we assumed equal sampling (15 sites along the cline). We compared the linear cline from the second comparison with a non-monotonic clines from the third comparison.

```
par(mar=c(4,4,3,0.4), mfrow=c(3,1))
plot(envPop_long$envPop, alFreq_L1_long, xlab="environment", ylab="allele freq.", col=cols[7], pch=pchs
abline(h=0.5, col=rgb(0,0,0,0.1))

plot(envPop_long$envPop, alFreq_nonMon, xlab="environment", ylab="allele freq.", col=cols[1], pch=pchs[
abline(h=0.5, col=rgb(0,0,0,0.1))

# CI vs. env
sequ <- seq(1, length(CI_nonMon), by=3)
plot(CI_nonMon[[1]]$x, CI_nonMon[[1]]$y, type="l", col=cols[1], ylim=c(-0.1,1), ylab="CI", xlab="enviro
for (i in sequ){
  points(CI_nonMon[[i]]$x, CI_nonMon[[i]]$y, type="l", col=adjustcolor(cols[1], 0.5))
  points(CI_long[[i]]$x, CI_long[[i]]$y, type="l", col=adjustcolor(cols[7], 0.5))
}

legend(-0.8, 1, fill=cols[c(7,1)], legend = c("cline", "non-mon."))
```

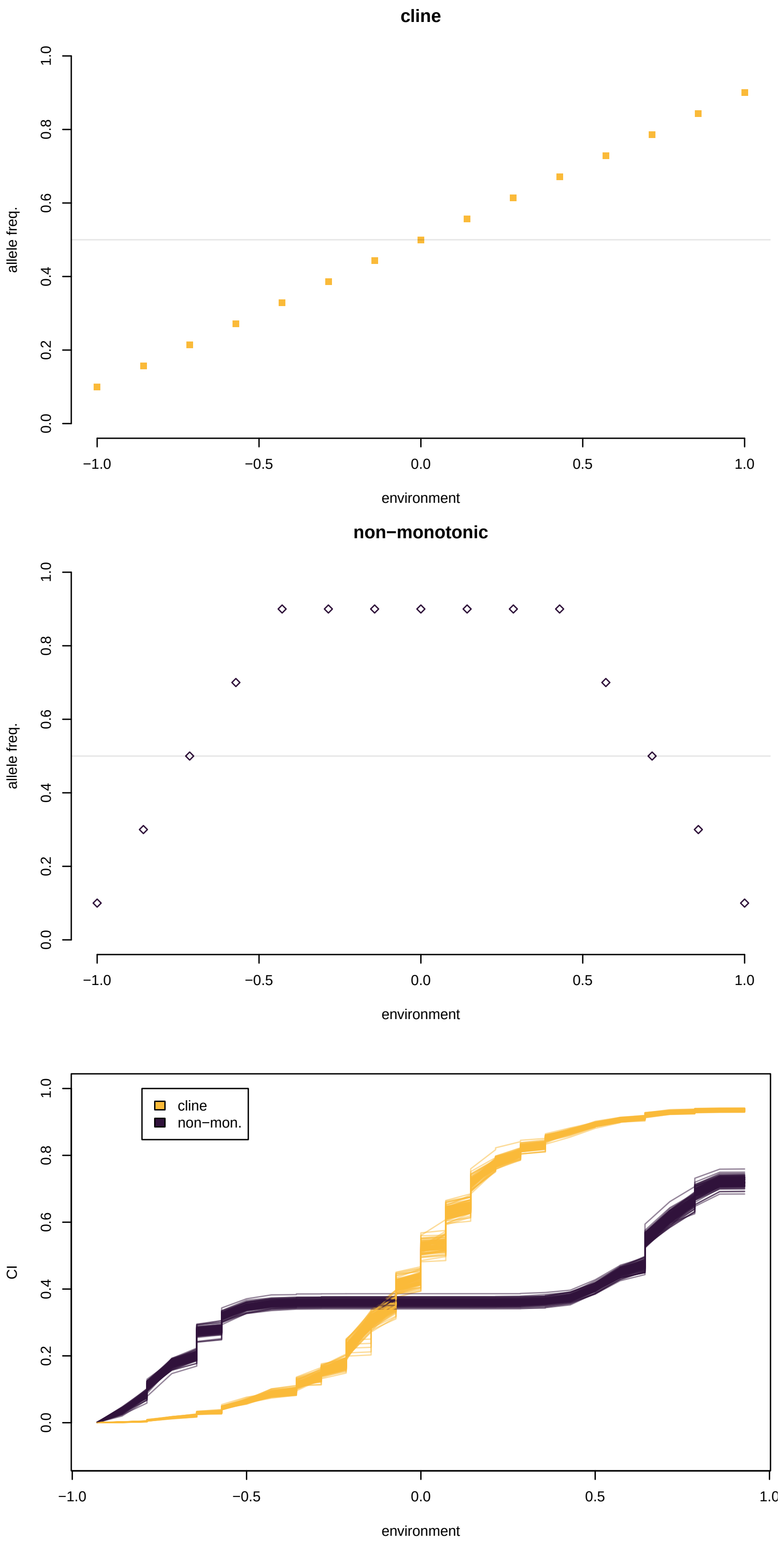
cline
allele freq.
environment
non−monotonic
allele freq.
environment
cline
non−mon.
CI
environment

The non-monotonic pattern has a flat CI in the intermediate values of the environment that have very little turnover. However, the clinal pattern, which is linear, results in a non-linear CI function. This illustrates a potential problem with interpreting the GF-offset as a genetic offset. For the cline at extreme values of the environment, a very small GF-offset would be calculated, and at intermediate values of the environment, a very large GF-offset would be calculated. However, the actual "genetic offset" (in terms of allele frequency change per unit environment change) is the same at all values of the environment.

### A steep cline vs. more randomness in the cline

#### Fifth take home message: The maximum value of the CI function reflects the amount of variation in the data that the model explains.

For this comparison, we assumed equal sampling (15 sites along the cline). We compared the linear cline from the second comparison with the same cline, but random noise added.

```
alFreq_L1_long_error <- alFreq_L1_long + rnorm(length(alFreq_L1_long), 0, 0.1)
alFreq_L1_long_error[alFreq_L1_long_error>1] <- 1
alFreq_L1_long_error[alFreq_L1_long_error<0] <- 0
CI_long_error <- replicate(100,runMod(envPop_long, alFreq_L1_long_error))
CI_long_error_R2 <- (unlist(CI_long_error[seq(3,length(CI_long_error), by=3)]))
```

```
par(mar=c(4,4,3,0.4), mfrow=c(3,1))
plot(envPop_long$envPop, alFreq_L1_long, xlab="environment", ylab="allele freq.", col=cols[7], pch=pchs
abline(h=0.5, col=rgb(0,0,0,0.1))

plot(envPop_long$envPop, alFreq_L1_long_error, xlab="environment", ylab="allele freq.", col=cols[3], pc
abline(h=0.5, col=rgb(0,0,0,0.1))

# CI vs. env
sequ <- seq(1, length(CI_long), by=3)
plot(CI_long[[1]]$x, CI_long[[1]]$y, type="l", col=cols[7], ylim=c(-0.1,1), ylab="CI", xlab="environmen
for (i in sequ){
  points(CI_long_error[[i]]$x, CI_long_error[[i]]$y, type="l", col=adjustcolor(cols[3], 0.5))
  points(CI_long[[i]]$x, CI_long[[i]]$y, type="l", col=adjustcolor(cols[7], 0.5))
}

legend(-0.8, 1, fill=cols[c(7,3)], legend = c("cline", "cline + noise"))
```

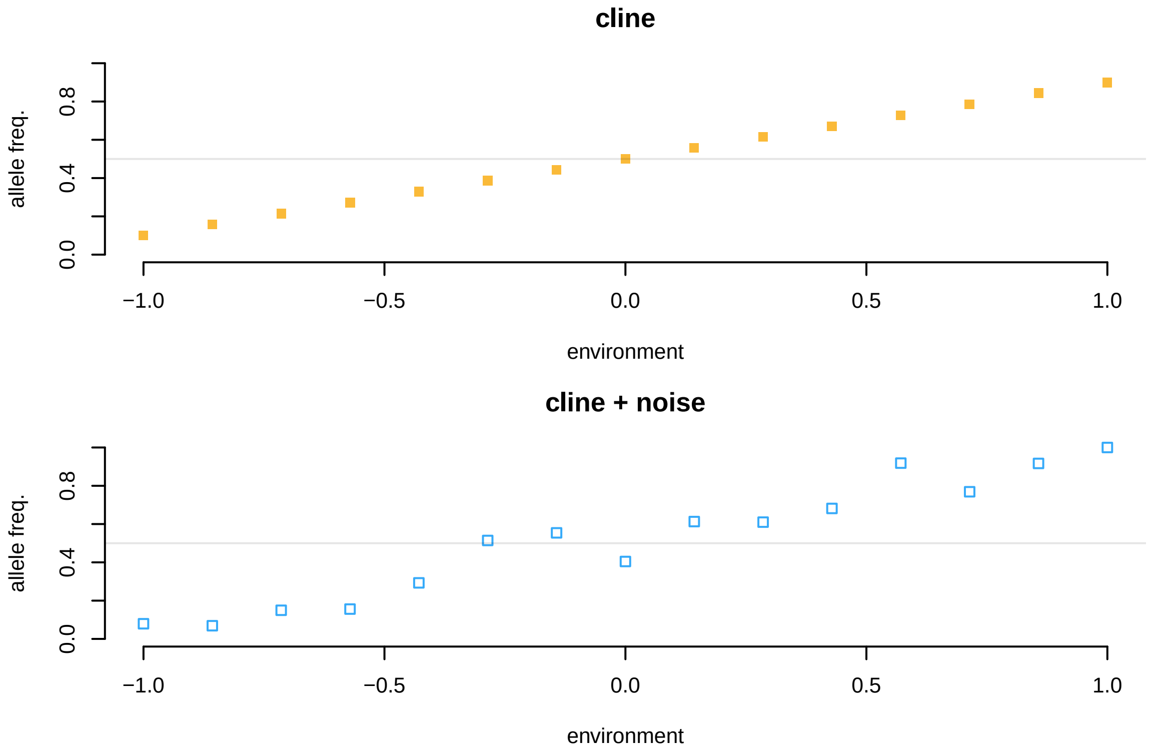


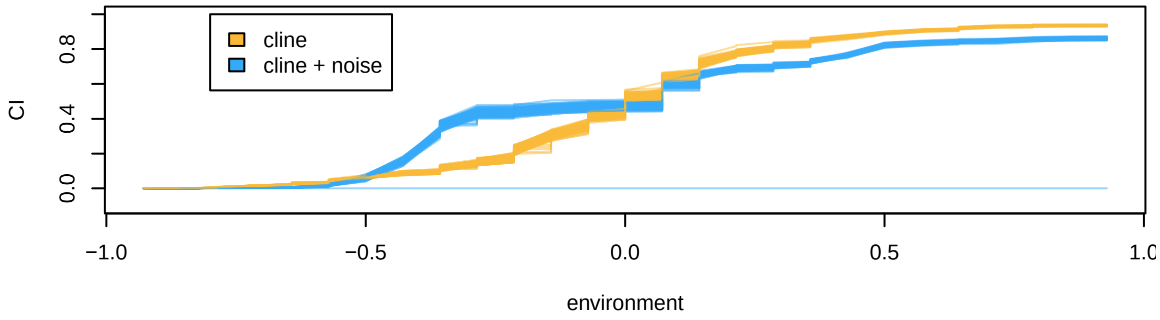


The above example shows that adding noise (as might occur due to genetic drift) changes the shape of the CI curve, and also lowers the maximum value that can be obtained. That is because the maximum value of the CI for an allele depends on how much variance in the allele frequency as a function of the environment the model can explain. When there is more noise in the data, the model explains less of the variance in allele frequency.

In addtion, we see the "all zeros in the CI despite high R^2" problem again with the noisy data, but it appeared to only happen once (and may not happen when the markdown renders).

```
sequ <- seq(1, length(CI_short), by=3)
plot(CI_long[[3]], max(CI_long[[1]]$y), col=adjustcolor(cols[7], 0.5), ylim=c(-0.1,1), xlab="R2", ylab=
for (i in sequ){
  points(CI_long[[i+2]], max(CI_long[[i]]$y), col=adjustcolor(cols[7], 0.5), pch=pchs[4])
  points(CI_long_error[[i+2]],max(CI_long_error[[i]]$y),  col=adjustcolor(cols[3], 0.5), pch=pchs[1])
}

legend(0.1, 1, col =cols[c(7,3)], legend = c("cline", "cline + error"), pch=pchs[c(4,1)])
```

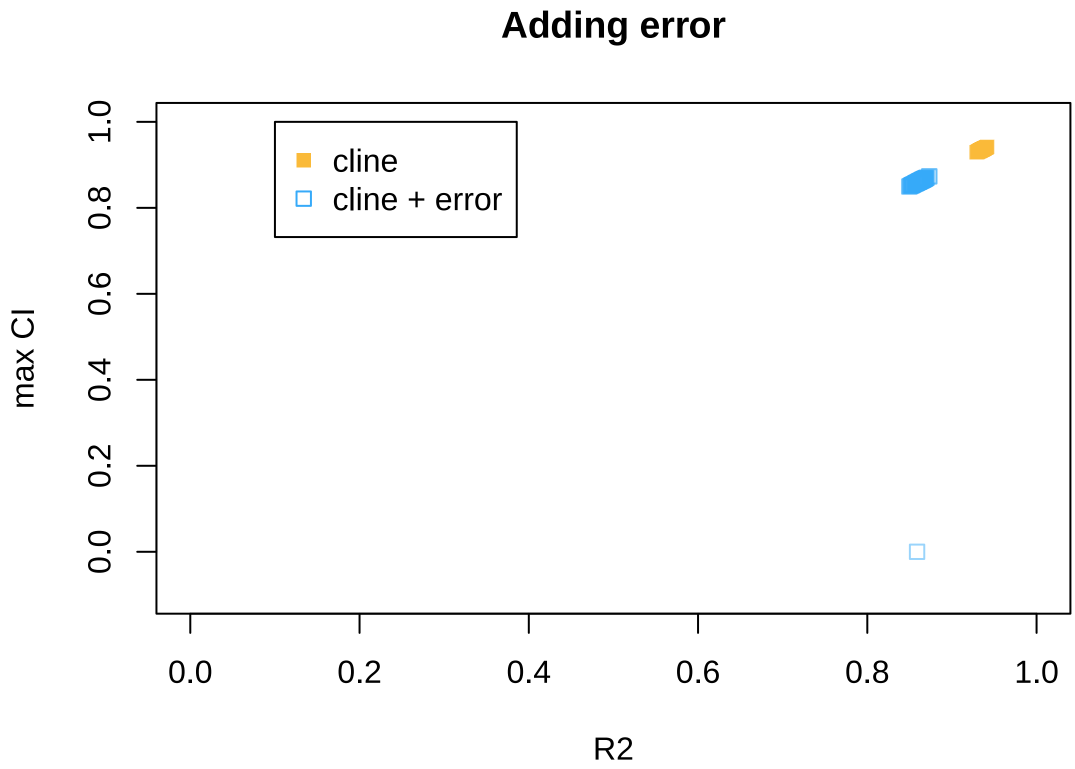


### Appendix: Varying the parameters

Here we examine: (1) varying number of trees and other parameters (2) varying number of loci and environments

##### testing smaller number of trees

The following chunck tests what happens when 50, instead of 500 trees are used in the gradient forest.

```
### 50 trees ####
envPop_short <- data.frame(envPop=seq(-1,1,length.out=6)) #create env
alFreq_L1_short<-seq(0.1,0.9,length.out=6)
CI_short <- replicate(100,runMod(envPop_short, alFreq_L1_short, 50))
(CI_short_prop0 <- sum(unlist(CI_short[seq(2,length(CI_short), by=3)]))/100)
```

```
## [1] 0.56
```

```
(CI_short_R2 <- (unlist(CI_short[seq(3,length(CI_short), by=3)])))
```

```
##    alFreq    alFreq    alFreq    alFreq    alFreq    alFreq    alFreq    alFreq
## 0.5961088 0.6156570 0.6020488 0.4997753 0.5847380 0.6595881 0.5129334 0.4943617
##    alFreq    alFreq    alFreq    alFreq    alFreq    alFreq    alFreq    alFreq
```

```
## 0.4792174 0.5998145 0.5636801 0.5915409 0.4756880 0.5949479 0.5647524 0.5135352
##    alFreq    alFreq    alFreq    alFreq    alFreq    alFreq    alFreq    alFreq
## 0.6427872 0.6243990 0.6051083 0.5930576 0.5941551 0.5377546 0.6118210 0.5357297
##    alFreq    alFreq    alFreq    alFreq    alFreq    alFreq    alFreq    alFreq
## 0.3302316 0.5904766 0.5978690 0.4600422 0.5469507 0.5459786 0.5204600 0.4988602
##    alFreq    alFreq    alFreq    alFreq    alFreq    alFreq    alFreq    alFreq
## 0.6187504 0.5036866 0.4419879 0.5143926 0.5710473 0.6234870 0.6943167 0.5814236
##    alFreq    alFreq    alFreq    alFreq    alFreq    alFreq    alFreq    alFreq
## 0.5138333 0.5412151 0.5271931 0.5409733 0.5341145 0.5462391 0.5968499 0.5338207
##    alFreq    alFreq    alFreq    alFreq    alFreq    alFreq    alFreq    alFreq
## 0.5053048 0.4823154 0.5430580 0.5769947 0.6491129 0.6024083 0.6568817 0.6632052
##    alFreq    alFreq    alFreq    alFreq    alFreq    alFreq    alFreq    alFreq
## 0.4730345 0.5986117 0.5543607 0.5980861 0.6160953 0.4533046 0.4791288 0.5850825
##    alFreq    alFreq    alFreq    alFreq    alFreq    alFreq    alFreq    alFreq
## 0.4839463 0.6189752 0.3779898 0.6734784 0.5509184 0.5787973 0.5584584 0.5204427
##    alFreq    alFreq    alFreq    alFreq    alFreq    alFreq    alFreq    alFreq
## 0.4719330 0.5846181 0.5884465 0.5197137 0.4723363 0.5870362 0.5681864 0.5695637
##    alFreq    alFreq    alFreq    alFreq    alFreq    alFreq    alFreq    alFreq
## 0.5415667 0.6097722 0.4813917 0.6471168 0.5380176 0.6199530 0.6352750 0.6247147
##    alFreq    alFreq    alFreq    alFreq    alFreq    alFreq    alFreq    alFreq
## 0.6377323 0.6141760 0.5135720 0.5115109 0.4513458 0.4735404 0.5449061 0.3705359
##    alFreq    alFreq    alFreq    alFreq
## 0.6807010 0.4461554 0.4385076 0.6126851
```

```
par(mfrow=c(1,1))
plot(CI_short[[3]], max(CI_short[[1]]$y), col=adjustcolor(cols[1], 0.5), ylim=c(-0.1,1), xlab="R2", yla
for (i in sequ){
  points(CI_short[[i+2]], max(CI_short[[i]]$y),  col=adjustcolor(cols[1], 0.5), pch=pchs[1])
}
```

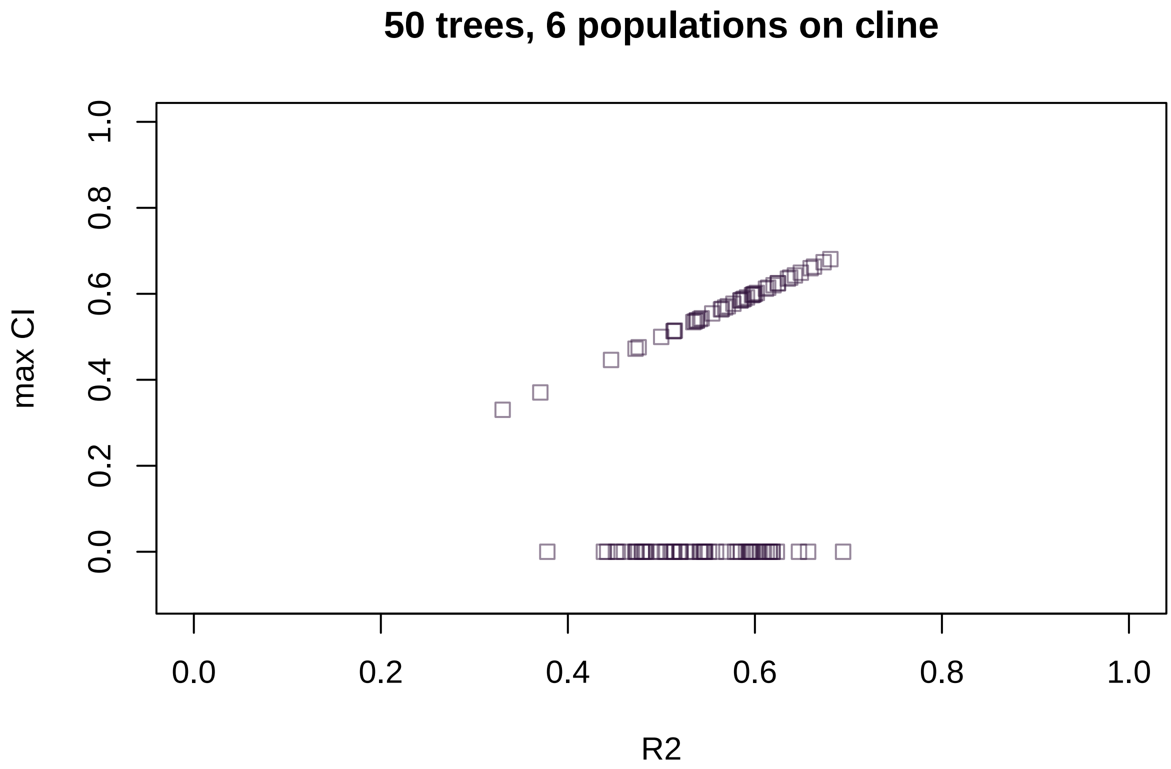


With 50 trees, there is higher variance in the R^2, but fewer replicates randomly have a max CI=0.

###### testing larger number of trees

The following chunck tests what happens when 3000, instead of 500 trees are used in the gradient forest.

```
### 3000 Trees
envPop_short <- data.frame(envPop=seq(-1,1,length.out=6)) #create env
alFreq_L1_short<-seq(0.1,0.9,length.out=6)
CI_short <- replicate(100,runMod(envPop_short, alFreq_L1_short, 3000))
(CI_short_prop0 <- sum(unlist(CI_short[seq(2,length(CI_short), by=3)]))/100)
```

```
## [1] 1
```

```
(CI_short_R2 <- (unlist(CI_short[seq(3,length(CI_short), by=3)])))
```

```
##    alFreq    alFreq    alFreq    alFreq    alFreq    alFreq    alFreq    alFreq
## 0.5756555 0.5781995 0.5815064 0.5710053 0.5809374 0.5725858 0.5744433 0.5804416
##    alFreq    alFreq    alFreq    alFreq    alFreq    alFreq    alFreq    alFreq
## 0.5838285 0.5709773 0.5789227 0.5776181 0.5817377 0.5577588 0.5834461 0.5841469
##    alFreq    alFreq    alFreq    alFreq    alFreq    alFreq    alFreq    alFreq
## 0.5694050 0.5774355 0.5847225 0.5808639 0.5793433 0.5835563 0.5862529 0.5697822
##    alFreq    alFreq    alFreq    alFreq    alFreq    alFreq    alFreq    alFreq
## 0.5727940 0.5847246 0.5841450 0.5927135 0.5842730 0.5864763 0.5920810 0.5913599
##    alFreq    alFreq    alFreq    alFreq    alFreq    alFreq    alFreq    alFreq
```

```
## 0.5932560 0.5788162 0.5785179 0.5870363 0.5855669 0.5978747 0.5925212 0.5921441
##    alFreq    alFreq    alFreq    alFreq    alFreq    alFreq    alFreq    alFreq
## 0.5817706 0.5842333 0.5819257 0.5794201 0.5940344 0.5791944 0.5941909 0.5909962
##    alFreq    alFreq    alFreq    alFreq    alFreq    alFreq    alFreq    alFreq
## 0.5944096 0.6038929 0.6018778 0.5699473 0.5972830 0.5886194 0.5809038 0.5893939
##    alFreq    alFreq    alFreq    alFreq    alFreq    alFreq    alFreq    alFreq
## 0.5800360 0.5867689 0.5921272 0.5859860 0.5911637 0.5674904 0.5931289 0.5884429
##    alFreq    alFreq    alFreq    alFreq    alFreq    alFreq    alFreq    alFreq
## 0.5949929 0.5757949 0.5748975 0.5779202 0.5926964 0.5840113 0.5745915 0.5767979
##    alFreq    alFreq    alFreq    alFreq    alFreq    alFreq    alFreq    alFreq
## 0.5876920 0.5780422 0.5862596 0.5818453 0.5787665 0.5756455 0.5763533 0.5839957
##    alFreq    alFreq    alFreq    alFreq    alFreq    alFreq    alFreq    alFreq
## 0.5797119 0.5896574 0.5846177 0.5815299 0.5731450 0.5824289 0.5809483 0.5814262
##    alFreq    alFreq    alFreq    alFreq    alFreq    alFreq    alFreq    alFreq
## 0.5729858 0.5999761 0.5913583 0.5803995 0.5856328 0.5935945 0.5764059 0.5893360
##    alFreq    alFreq    alFreq    alFreq
## 0.5837130 0.5736573 0.5771753 0.5794884
```

```r
par(mfrow=c(1,1))
plot(CI_short[[3]], max(CI_short[[1]]$y), col=adjustcolor(cols[1], 0.5), ylim=c(-0.1,1), xlab="R2", yla
for (i in sequ){
  points(CI_short[[i+2]], max(CI_short[[i]]$y),  col=adjustcolor(cols[1], 0.5), pch=pchs[1])
}
```

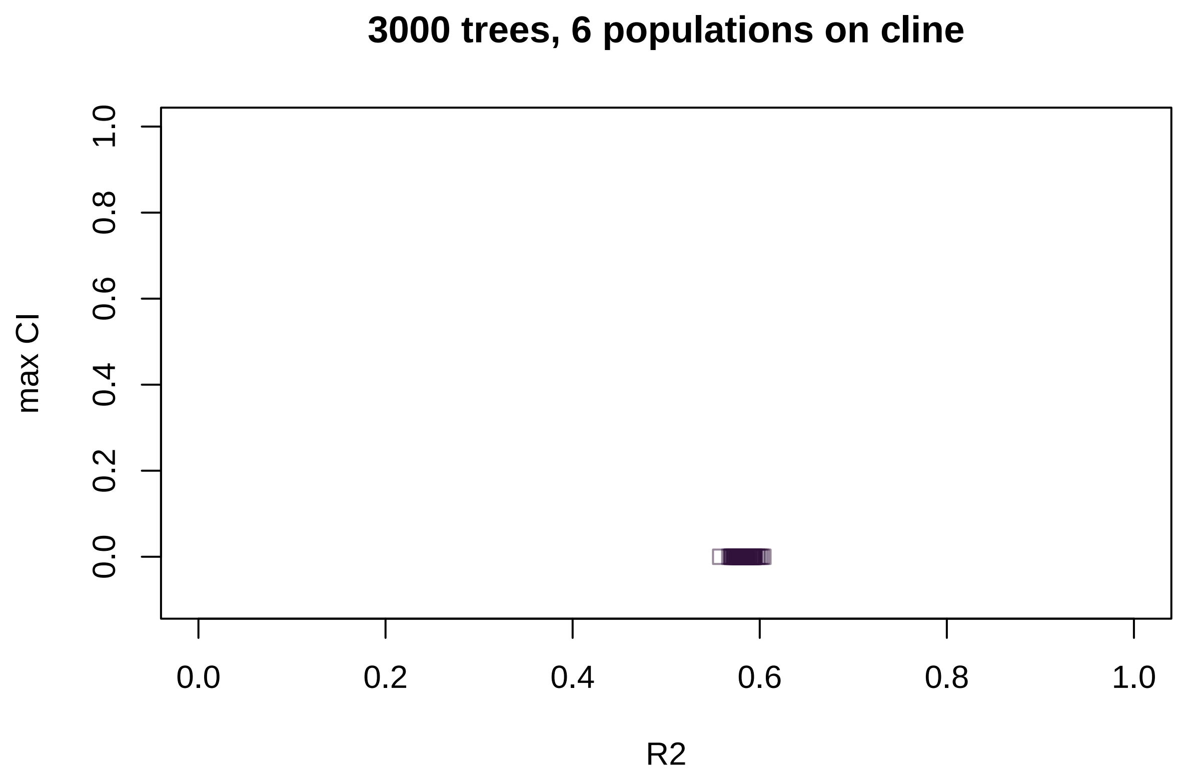


With 3000 trees, there is lower variance in the R^2, but all replicates have a max CI=0. That was weird. Moving on.

###### testing higher corr threshold

The following chunck tests if different correlation thresholds in the gradient forest affects the output.

```
### Compare the same cline for 10 vs. 50 environment sites ####
envPop_short <- data.frame(envPop=seq(-1,1,length.out=8)) #create env
alFreq_L1_short<-seq(0.1,0.9,length.out=8)
CI_short <- replicate(100,runMod(envPop_short, alFreq_L1_short, 500, 0.9))
print("Proportion of times we get all 0's in the CI with 8 sites, 500 trees, cor threshold 0.9:")
```

```
## [1] "Proportion of times we get all 0's in the CI with 8 sites, 500 trees, cor threshold 0.9:"
```

```
(CI_short_prop0 <- sum(unlist(CI_short[seq(2,length(CI_short), by=3)]))/100)
```

```
## [1] 0.7
```

```
CI_short <- replicate(100,runMod(envPop_short, alFreq_L1_short, 500, 0.5))
print("Proportion of times we get all 0's in the CI with 8 sites, 500 trees, cor threshold 0.5:")
```

```
## [1] "Proportion of times we get all 0's in the CI with 8 sites, 500 trees, cor threshold 0.5:"
```

```
(CI_short_prop0 <- sum(unlist(CI_short[seq(2,length(CI_short), by=3)]))/100)
```

```
## [1] 0.69
```

```
CI_short <- replicate(100,runMod(envPop_short, alFreq_L1_short, 500, 0.1))
print("Proportion of times we get all 0's in the CI with 8 sites, 500 trees, cor threshold 0.1:")
```

```
## [1] "Proportion of times we get all 0's in the CI with 8 sites, 500 trees, cor threshold 0.1:"
```

```
(CI_short_prop0 <- sum(unlist(CI_short[seq(2,length(CI_short), by=3)]))/100)
```

```
## [1] 0.59
```

```
CI_short <- replicate(100,runMod(envPop_short, alFreq_L1_short, 50, 0.1))
print("Proportion of times we get all 0's in the CI with 8 sites, 50 trees, cor threshold 0.1:")
```

```
## [1] "Proportion of times we get all 0's in the CI with 8 sites, 50 trees, cor threshold 0.1:"
```

```
(CI_short_prop0 <- sum(unlist(CI_short[seq(2,length(CI_short), by=3)]))/100)
```

```
## [1] 0.07
```

```
CI_short <- replicate(100,runMod(envPop_short, alFreq_L1_short, 1000, 0.1))
print("Proportion of times we get all 0's in the CI with 8 sites, 1000 trees, cor threshold 0.1:")
```

```
## [1] "Proportion of times we get all 0's in the CI with 8 sites, 1000 trees, cor threshold 0.1:"
```

```r
(CI_short_prop0 <- sum(unlist(CI_short[seq(2,length(CI_short), by=3)]))/100)
```

```
## [1] 0.87
```

```r
CI_short <- replicate(100,runMod(envPop_short, alFreq_L1_short, 10, 0.1))
print("Proportion of times we get all 0's in the CI with 8 sites, 10 trees, cor threshold 0.1:")
```

```
## [1] "Proportion of times we get all 0's in the CI with 8 sites, 10 trees, cor threshold 0.1:"
```

```r
(CI_short_prop0 <- sum(unlist(CI_short[seq(2,length(CI_short), by=3)]))/100)
```

```
## [1] 0.03
```

```r
(CI_short_R2 <- (unlist(CI_short[seq(3,length(CI_short), by=3)])))
```

```
##    alFreq    alFreq    alFreq    alFreq    alFreq    alFreq    alFreq    alFreq
## 0.6368457 0.7317672 0.8094167 0.6500869 0.7224698 0.7363063 0.7033746 0.4337905
##    alFreq    alFreq    alFreq    alFreq    alFreq    alFreq    alFreq    alFreq
## 0.6619707 0.7372738 0.5977575 0.5011033 0.5453371 0.2007081 0.7694146 0.6684384
##    alFreq    alFreq    alFreq    alFreq    alFreq    alFreq    alFreq    alFreq
## 0.7326787 0.7181097 0.7718453 0.6321164 0.6140487 0.6170833 0.6578323 0.5858349
##    alFreq    alFreq    alFreq    alFreq    alFreq    alFreq    alFreq    alFreq
## 0.5742056 0.7559343 0.6010065 0.7191471 0.6135152 0.5126286 0.5536318 0.5035281
##    alFreq    alFreq    alFreq    alFreq    alFreq    alFreq    alFreq    alFreq
## 0.4395559 0.6597501 0.7837763 0.6345325 0.6126654 0.7652169 0.6185079 0.4293853
##    alFreq    alFreq    alFreq    alFreq    alFreq    alFreq    alFreq    alFreq
## 0.5856088 0.6378630 0.7123296 0.6008612 0.5576720 0.6789379 0.6992388 0.7771136
##    alFreq    alFreq    alFreq    alFreq    alFreq    alFreq    alFreq    alFreq
## 0.6260177 0.3707786 0.6879255 0.6710652 0.6799350 0.1496778 0.5628693 0.5162126
##    alFreq    alFreq    alFreq    alFreq    alFreq    alFreq    alFreq    alFreq
## 0.5457196 0.6678610 0.7155342 0.5425138 0.6788277 0.6504144 0.3545991 0.4547625
##    alFreq    alFreq    alFreq    alFreq    alFreq    alFreq    alFreq    alFreq
## 0.6662693 0.5672082 0.6964964 0.3885731 0.7005338 0.3956479 0.5215172 0.7549485
##    alFreq    alFreq    alFreq    alFreq    alFreq    alFreq    alFreq    alFreq
## 0.6444631 0.7441090 0.6749391 0.7108058 0.4340255 0.6356566 0.7502808 0.7142720
##    alFreq    alFreq    alFreq    alFreq    alFreq    alFreq    alFreq    alFreq
## 0.4959523 0.6473906 0.7126496 0.6666058 0.7396416 0.5685130 0.5828108 0.4957539
##    alFreq    alFreq    alFreq    alFreq    alFreq    alFreq    alFreq    alFreq
## 0.6222979 0.6706362 0.4801448 0.5864618 0.4320888 0.4171146 0.7256306 0.6648753
##    alFreq    alFreq    alFreq    alFreq
## 0.4450046 0.7322682 0.4858871 0.6562764
```

```r
par(mfrow=c(1,1))
plot(CI_short[[3]], max(CI_short[[1]]$y), col=adjustcolor(cols[1], 0.5), ylim=c(-0.1,1), xlab="R2", ylal
for (i in sequ){
  points(CI_short[[i+2]], max(CI_short[[i]]$y),  col=adjustcolor(cols[1], 0.5), pch=pchs[1])
}
```

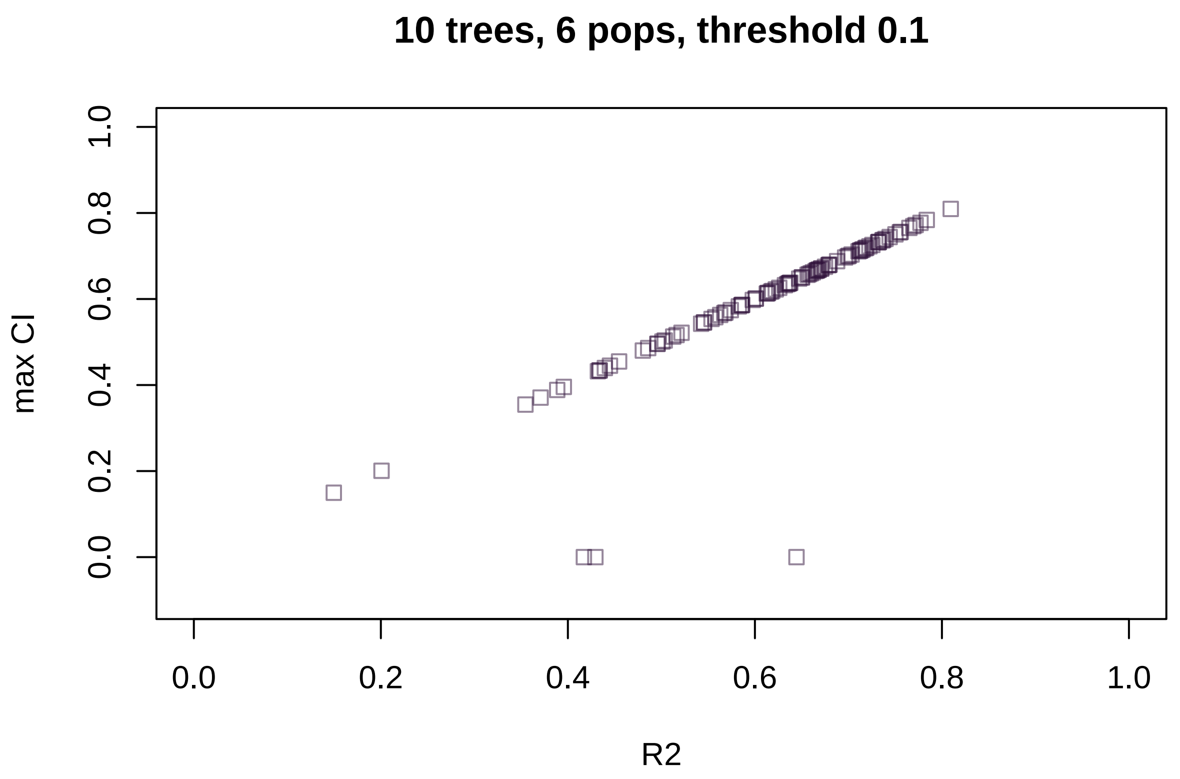


I'm wondering if this has something to do with the number of trees far exceeding the number of possible splits given only a few samples along a gradient. When the number of requested permutations exceeds the number possible in the data, the result is a large number of identical trees (?).

I did find that I was getting errors when the number of tress was *less* then the number of samples in the data.

###### testing multiple environmental predictors

Here, I compare two models. The first "univariate" model only includes the true environmental predictor. The second "multivariate model" includes the true predictor and two random environmental variables.

Each has 15 sites sampled along the true environmental gradient.

```
envPop_short <- data.frame(envPop=seq(-1,1,length.out=15),
                           env2 = rnorm(15),
                           env3 = rnorm(15)) #create env
alFreq_L1_short<-seq(0.1,0.9,length.out=15)

uni <- runMod(data.frame(envPop=envPop_short[,1]), alFreq_L1_short)
multi <- runMod_multi(envPop_short, alFreq_L1_short)

print("The univariate model, which only includes the true predictor, has an R^2 of:")
print(uni$R2)

print("The multivariate model, which includes the true predictor and two random predictors, has an R^2 o
```

```
print(multi$R2)

# How are the CI functions affected by including multiple predictors?
plot(uni$CI$x, uni$CI$y, main="compare univariate (black circ.)\nto multivariate (blue tri.)")
points(multi$CI_env$envPop$x, multi$CI_env$envPop$y, col="blue", pch=2)
```

The above R^2 results make sense. When more random predictors are included, the random forest would choose these sometimes to build the model, and this would lower the overal predictive power of the model compared to a model that only includes the true predictor.

The plot shows that when adding additional random environmental predictors, it affects the total CI of the model for the true environmental predictor. In addition this would affect genetic offset calculations near the environmental extremes.

##### Testing multiple loci and one environment

Here we ask, does adding more loci to the model affect the results for one locus? Based on first princples, it should not, because each locus is an independent model. I created a second allele with frequencies sampled from a beta distribution, and a third allele with frequencies sampled randomly from the first locus. The second allele does not have a positive R^2 in a univariate model, but the third allele does. Here, I test if the statistics output from the 1-locus-1-environment model are the same as the multi-locus-multi-environment model.

```
L1 = seq(0.1,0.9,length.out=15)
L2 = rbeta(15, 1, 1)
set.seed(9126)
L3 = sample(L1)
envPop_short <- data.frame(envPop=seq(-1,1,length.out=15)) #create env
alFreq_L1_short<-data.frame(L1,L2, L3)
# use "uni" model above
uni
# uni_L2 <- runMod(envPop_short, L2) this returns NULL
uni_L3 <- runMod(envPop_short, L3)
uni_L3
uni_L3_2 <- runMod(envPop_short, L3)
uni_L3_2

multi_loc <- runMod_multi(envPop_short, alFreq_L1_short)
multi_loc

plot(uni$CI$y, multi_loc[[5]]$envPop$L1$y, xlab="CI L1 one locus", ylab="CI L1 three locus")
abline(0,1)

plot(uni_L3$CI$y, uni_L3_2$CI$y, xlab="CI L3 one locus", ylab="CI L3 one locus again")
abline(0,1)

plot(uni_L3$CI$y, multi_loc[[5]]$envPop$L3$y[2:43], xlab="CI L3 one locus", ylab="CI L3 three locus")
abline(0,1)

print("The R^2 for L1 from the one-locus model:")
print(uni$R2)
print("The R^2 for L1 from the multi-locus model:")
print(multi_loc$R2[1])
```

```
print("The R^2 for L3 from the one-locus model:")
print(uni_L3$R2)
print("The R^2 for L3 from the multi-locus model:")
print(multi_loc$R2[2])
```

The last analysis shows that adding more loci does not affect the CI curves for a single locus, above and beyond what happens from run to run given the randomness in the forest.
